# Metacommunities, fitness and gradual evolution

**DOI:** 10.1101/2021.01.15.426821

**Authors:** Tadeas Priklopil, Laurent Lehmann

## Abstract

We analyze the evolution of a multidimensional quantitative trait in a class structured focal species interacting with other species in a wider metacommunity. The evolutionary dynamics in the focal species as well as the ecological dynamics of the whole metacommunity is described as a continuous time process with birth, physiological development, dispersal, and death given as rates that can depend on the state of the whole metacommunity. This can accommodate complex local community and global metacommunity environmental feedbacks owing to inter- and intra-specific interactions, as well as local environmental stochastic fluctuations. For the focal species, we derive a fitness measure for a mutant allele affecting class-specific trait expression. Using classical results from geometric singular perturbation theory, we provide a detailed proof that if the effect of the mutation on phenotypic expression is small (“weak selection”), the large system of dynamical equations needed to describe selection on the mutant allele in the metacommunity can be reduced to a single ordinary differential equation on the arithmetic mean mutant allele frequency that is of constant sign. This invariance on allele frequency entails the mutant either dies out or will out-compete the ancestral resident (or wild) type. Moreover, the directional selection coefficient driving arithmetic mean allele frequency can be expressed as an inclusive fitness effect calculated from the resident metacommunity alone, and depends, as expected, on individual fitness differentials, relatedness, and reproductive values. This formalizes the Darwinian process of gradual evolution driven by random mutation and natural selection in spatially and physiologically class structured metacommunities.

---

> Invariance that had appeared in the criterion for altruism with respect to gene frequency in the case of sibs had seemed a gift from God, and I did not expect to see it repeated in the more complex trial cases I had moved on to. So it was with joy and almost with incredulity that I at last found emerging out of acres of my tedious and usually wrong algebra for the case of uncles, and then for the case of cousins, the same invariance as I had found before.
>
> — (Hamilton, 1988, p. 189)

## 1 Introduction

Darwinian evolution – the gradual, step-by-step transformation of traits due to random mutation and non-random cumulative natural selection – is the central mechanism of adaptation (Dawkins, 1986, 1997). A proof of principle of Darwinian evolution is given by selective sweeps, in that mutations that increase in frequency also tend to fix in the population as a result of natural selection. Each fixation thus displays invariance in the direction of selection with respect to gene frequency, and repeated fixations under the constant influx of mutations enable gradual phenotypic evolution and adaptation. Such invariance with respect to gene frequency is, however, not expected to hold under general biological processes. In particular, social as well as ecological interactions can lead to survival and reproduction being frequency-dependent, and this may result in genetic polymorphisms preventing allelic fixation. Furthermore, in spatially and physiologically structured populations, it is not obvious to identify the conditions under which selective sweeps occur owing to the complications arising from genetic correlations at local spatial scales and across physiological classes.

In this paper we provide sufficient conditions for selective sweeps when evolution occurs in an asexual focal species that resides in a wider metacommunity (Wilson, 1992; Leibold et al., 2004), and where socially interacting individuals are structured into discrete classes that determine their physiological state, such as age and size, as well as their local biotic and abiotic environment. We provide these conditions in terms of an “invasion implies substitution”-principle, which is a formal characterization of the invariance of selection with respect to a population-wide frequency of a mutant allele that affects quantitative traits in the context of density- and frequency-dependent selection (Hamilton, 1964; Rousset, 2004; Meszéna et al., 2005; Dercole and Rinaldi, 2008; Lehmann and Rousset, 2014; Dercole, 2016; Cantrell et al., 2017; Ito et al., 2020; Cai and Geritz, 2020; Priklopil and Lehmann, 2020). The principle consists of two propositions (Priklopil and Lehmann, 2020). The first says that if a mutation in the focal species results in a phenotype with a closely similar expression to its ancestral resident (wild)-type, a successful invasion of the mutant allele will generically lead to its fixation. In the present metacommunity model, moreover, the dynamical process for some weighted population-wide mutant frequency in the focal species *p*(*t*) ∈ [0, 1] is supposed to take the form of a scalar-valued ordinary differential equation

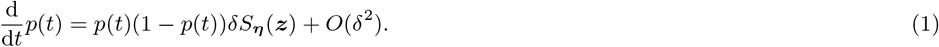

Here, *δ* is a small parameter characterizing the distance between the mutant phenotype ***z*** + *δ****η*** and the resident phenotype ***z*** = (*z*_A_)_A∈𝒞_, with *z*_A_ denoting the phenotype produced by a carrier of the resident allele in class A ∈ 𝒞 where 𝒞 is the set of relevant classes individuals can be in, and where ***η*** = (*η*_A_)_A∈𝒞_ is the mutant phenotypic deviation. The expression *δS*_***η***_ (***z***) in (1) is the so-called directional selection coefficient, and because it is frequency-independent the allele frequency affects the evolutionary dynamics only via the term *p*(*t*)(1 – *p*(*t*)). Since *S*_***η***_ (***z***) is independent of time *t*, a successful invasion of the mutant allele implies its substitution and thus a selective sweep is obtained in the focal species. This in turn implies that in longer timescales recurrent mutation and invasion events will gradually transform the phenotype ***z*** of interest until the phenotype reaches the boundary of its feasible values or *S*_***η***_ (***z***) becomes zero, which allows to identify singular points where the selection on ***z*** changes from directional to either stabilizing or disruptive (e.g. Geritz et al., 1998; Rousset, 2004).

The second proposition says that the rate of substitution of the invading mutant allele is model independent, and that it can be expressed in terms of an inclusive fitness effect as

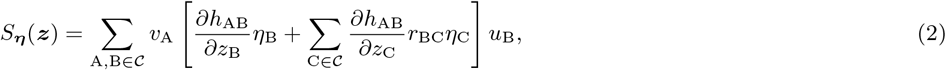

where all quantities are calculated from the resident population. Here, *h*_BA_ is an element of a resident growth-rate matrix **H** giving the rate at which a single individual in class A ∈ 𝒞 that expresses phenotype *z*_A_ produces individuals in class B ∈ 𝒞. The first partial derivative inside the brackets is taken with respect to the phenotype of the focal individual whose growth-rate we are considering and measures the (direct fitness) effect of the mutation on its own growth-rate, where the mutational effect is of magnitude *η*_B_. The second partial derivative is taken with respect to the phenotype of an individual of class C inhabiting the same group as the ‘focal’ individual, and measures the (indirect fitness) effect that mutant individuals in class C have on the growth-rate of the focal individual. This derivative is weighted with the relatedness coefficient *r*_BC_ giving the probability that an individual in class C ∈ 𝒞 and the focal individual in class B ∈ 𝒞 have a common ancestor, that is, their lineages coalesce backward in time. The vector **r** = (*r*_BC_)_B,C∈𝒞_ collecting all these relatedness coefficients thus gives the conditional probability of group members inheriting the same allele, and captures the effect of limited genetic mixing on the evolutionary dynamics (**r** = **0** in spatially well-mixed species). Finally, the vectors **u** = (*u*_A_)_A∈𝒞_ and **v** = (*v*_A_)_A∈𝒞_ are the right and left dominant eigenvectors of **H** and are normalized to give class frequencies and reproductive values, respectively. Expressions taking broadly the form (2) and special cases thereof are routinely applied in evolutionary biology to ascertain the directions of selection and evaluate singular points, be it in the kin selection, quantitative genetics, life-history, demography, and adaptive dynamics literatures (e.g., Charlesworth, 1994; Frank, 1998; Caswell, 2000; Rousset, 2004; Dercole and Rinaldi, 2008; Lynch and Walsh, 2018 for textbook treatments).

In this paper, we extend our earlier result on the “invasion implies substitution”-principle (1)-(2) derived for well-mixed ecological communities (Priklopil and Lehmann, 2020) to spatially structured metacommunities where local communities are of finite size and follow an island model of dispersal. For such spatial structure, the “invasion implies substitution”-principle has been previously considered in Roze and Rousset (2003, 2004); Rousset and Ronce (2004); Rousset (2004); Van Cleve et al. (2010) [see also closely related results in Rousset, 2006; Lion and Gandon, 2009 for lattice spatial structure and Cantrell et al., 2017 where dispersal follows a diffusion model], and our approach and results here extend and complement this earlier work in various aspects. First, we formulate our model in continuous time and for several interacting and stochastically reproducing species that are physiologically class structured. Second, we provide a step-by-step proof of (1)-(2) by singularly perturbing the limiting metacommunity model where *δ* = 0 (Fenichel, 1979; Hek, 2010; Kuehn, 2015; Priklopil and Lehmann, 2020). Third, we pay particular attention to tie up all underlying concepts needed to prove gradual evolution in metacommunities, from the individual based demographic processes to the dynamics of moments of allelic states and the fast-slow analysis of all the relevant dynamical variables. We find that the approximation (1)-(2) holds for small but non-zero *δ* where the perturbation of steady states caused by the invasion is taken into account, a result given without proof previously. More importantly, we sharpen earlier representations of (1) in terms of the reproductive value weighted average frequency (Roze and Rousset, 2004; Rousset and Ronce, 2004; Rousset, 2004; Van Cleve et al., 2010; Priklopil and Lehmann, 2020) by allowing *p*(*t*) to be defined as any population-wide average mutant frequency, with special cases being the reproductive value weighted mutant frequency as well as the arithmetic mean mutant frequency. In terms of the arithmetic mean, equation (1) becomes functionally equivalent to the standard and widely applied textbook representation of natural selection under additive gene action in spatially and physiologically homogeneous populations without social interactions nor ecological feedback (e.g., Crow and Kimura, 1970; Gillespie, 2004; Hartl et al., 1997).

The rest of the paper is organized as follows. In Section 2 we introduce the main modeling assumptions and in Section 3 we detail the specifics of the purely resident (ecological) metacommunity model. All the species dynamical variables and growth-rates that are present in the directional selection coefficient *δS*_***η***_ (**z**) as characterised in (2) are derived in this section. The remainder of the paper is then dedicated in proving the “invasion implies substitution”-principle (1)-(2) for the metacommunity model. To do this, we first present in Section 4 a mutant-resident metacommunity model for arbitrary mutant and resident phenotypes and thus arbitrary strength of selection. Then, in Sections 5-6, we will derive several results for the limiting metacommunity processes where *δ* = 0, and the formal proof of the “invasion implies substitution”-principle where *δ* is small but non-zero is given in Section 7. In Section 8 we will conclude the paper by discussing the results and a particular focus is given in relating our results to the theory of adaptation and gradual evolution.

## 2 Outline of the model

We consider a metacommunity consisting of an infinite number of local communities connected to each other by random uniform dispersal. Each local community, which we will generally refer to as a local group, consists of a finite number of interacting asexual species and we assume each local community to be bounded in size. We focus on the ecological and evolutionary dynamics occurring in some focal species of interest and assume a continuous time demographic process.

At each moment in time, each individual in the focal species is assumed to be in one of a finite number *k*_𝒫_ of individual-level (i-level) physiological states, such as of certain age and size, and we denote the set of physiological states of the focal species with 𝒫. All the other non-focal species can be similarly structured, and in addition each group is characterized by a set of *abiotic factors* such as the temperature or quality of the habitat. An abiotic factor is defined as any factor outside of the focal and non-focal species, be it a factor that is affected by the species or not, in which case we call it an external factor. Each group of individuals will therefore be in one of a finite number *k*_𝒮_ of group-level states, defined as a union of physiological states of each and every individual of every species inhabiting the group, including any abiotic factors. The set of all group states (g-state) is denoted with 𝒮. As each element of 𝒮 prescribes exactly the size distributions of all species, including the focal, we will use *N*_*as*_ to denote the number of individuals *a* ∈ 𝒫 in the focal species inhabiting a group in g-state *s* ∈ 𝒮.

Each group in the metacommunity changes its state in 𝒮 due to individual processes (i-processes) of the focal and non-focal species, as well as due to processes of abiotic factors such as changes in the temperature or quality of the habitat. The i-processes of the focal species are birth, where offspring of any physiological state is being produced, and dispersal, where each offspring either disperses randomly to other groups or stays in its natal group. Individuals may also undergo physiological development defined as a transition from one physiological state to another, as well as death. Whenever a group *s* ∈ 𝒮 transitions to *u* ∈ 𝒮 due to an event where an individual *a* ∈ 𝒫 is produced into or removed from the group, we have the identities *N*_*as*_ + 1 = *N*_*au*_ or *N*_*as*_ – 1 = *N*_*au*_, respectively, and otherwise *N*_*as*_ = *N*_*au*_. Individuals of the other non-focal species may undergo similar i-process. All the processes are assumed stochastic occurring on a Poissonian basis (e.g., Kingman, 1992), and can depend in an arbitrary way on the physiological state of the individual, the g-state of the group it inhabits as well as the state of all the other non-focal groups.

Because the i-procesess of individuals may differ not only due to their physiological state but also the g-state of the group they inhabit, we will use the notion of a *class* of an individual. This is defined such that whenever any two individuals are identical in terms of their i-processes, they belong to the same class (Taylor, 1990; Rousset, 2004; Grafen, 2015). In the present model, individuals are in the same class whenever they are in the same physiological state *and* inhabit a group that is in the same g-state. The class space is denoted with 𝒞 = 𝒫 × 𝒮 and it is of size *k*_𝒞_ = *k*_𝒫_ × *k*_𝒮_.

Finally, individuals in the focal species express a genetically evolving phenotype that may affect the i-processes of any individual in the metacommunity. We assume that phenotypic expression is a class-specific function. For example, the phenotype may depend on the physiological state of the individual as well as the abiotic factors of the group that it inhabits. An individual in class *as* ∈ 𝒞 expresses phenotype *z*_*as*_ ∈ ℝ such that ***z*** = (*z*_*as*_)_*as*∈𝒞_ ∈ *Ƶ* denotes the vector-valued phenotypic profile across classes and where 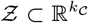 is the phenotype space. We assume that offspring inherit faithfully the phenotype of their parent and that two alleles that can result in different phenotypic expression segregate in the focal species, a mutant allele M and a resident allele R. We assume that the mutation rate is exceedingly small so that no other alleles interfere with the mutant-resident dynamics (this assumption can be mildly relaxed, see Section 8). Characterizing the spread of the mutant allele is the focus of our analysis.

We note that many closely connected ecological or evolutionary models allowing for various heterogeneities within and/or between groups have been considered before for both continuous or discrete time processes e.g. by Chesson (1981, 1985); Grey et al. (1995); Frank (1998); Gandon and Michalakis (1999); Ronce et al. (2000); Metz and Gyllenberg (2001); Arrigoni (2003); Cadet et al. (2003); Barbour and Pugliese (2004); Rousset (2004); Rousset and Ronce (2004); Martcheva and Thieme (2005); Lehmann et al. (2006); Martcheva and Thieme (2006); Martcheva et al. (2006); Alizon and Taylor (2008); Wild et al. (2009); Ronce and Promislow (2010); Wild (2011); Peña (2011); Rodrigues and Gardner (2012); Parvinen (2013); Massol and Débarre (2015); Lehmann et al. (2016); Rodrigues (2018); Parvinen et al. (2018); Kuijper and Johnstone (2019); Ohtsuki et al. (2020). No prior model has however considered metacommunities with multiple interacting species where individuals and groups are characterized by arbitrary states and as such all these previous models, as well as models with finite class-structure in panmictic populations, can be conceptually subsumed to the present analysis.

## 3 Resident dynamics in the metacommunity

The aim here is to describe the population dynamics (ecology) of the focal species in the metacommunity assuming that the focal species is monomorphic for the resident allele. For this, we need to characterize the dynamics of the entire resident metacommunity. We define the state of the resident monomorphic metacommunity as a discrete probability (frequency) distribution over the state space of groups 𝒮. The metacommunity state space is therefore the space of frequency distributions Δ(𝒮) on 𝒮, which is the simplex in 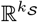. We will represent the frequency distribution of the metacommunity at time *t* ∈ ℝ with the vector **x** := **x**(*t*), where **x** = (*x*_*s*_)_*s*∈𝒮_ ∈ Δ(𝒮) and where *x*_*s*_ is the frequency of groups in state *s* ∈ 𝒮 with ∑ _*s*∈𝒮_ *x*_*s*_ = 1, and next we track the dynamics of this vector. While the upcoming characterization may at first appear lengthy, we fully work it out as (i) it plays a central role in the analysis of selection and (ii) it is of interest in the formulation of purely metacommunity ecological models.

### 3.1 Individual-level processes

To construct dynamical equations for the metacommunity dynamics, we start from i-processes of the focal species. To that end, we introduce four elementary individual-level rates (i-rates). First, the rate *γ*_*a*←*bs*_ at which an individual in physiological state *b* ∈ 𝒫 inhabiting a group *s* ∈ 𝒮, that is, an individual in class *bs* ∈ 𝒞, transitions into physiological state *a* ≠ *b* ∈ 𝒫. By construction, ∑ _*a*≠*b*∈𝒫_ *γ*_*a*←*bs*_ then stands for the total rate at which an individual in class *bs* ∈ 𝒞 undergoes a transition and *γ*_*a*←*bs*_ is undefined as a transition rate for *a* = *b* ∈ 𝒫. Second, the rate *µ*_*bs*_ at which an individual in class *bs* ∈ 𝒞 dies. Third, the rate 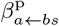 at which an individual in class *bs* ∈ 𝒞 produces by reproduction an offspring individual in physiological state *a* ∈ 𝒫 that stays in the focal group (hence superscript ‘p’ for philopatry), and finally, the rate 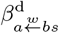 at which an individual in class *bs* ∈ 𝒞 produces by reproduction an offspring individual *a* ∈ 𝒫 into a non-focal group *w* ∈ 𝒮, conditional on offspring landing in that group (hence superscript ‘d’ for dispersal). We further assume that dispersed offspring land into a group *w* ∈ 𝒮 with probability *x*_*w*_, and this assumption allows us to write 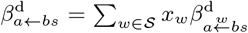 giving the total rate at which *bs* ∈ 𝒞 gives birth to *a* ∈ 𝒫 into non-focal groups (Figure 1). By convention we assume that whenever *N*_*bs*_ = 0 then *γ*_*a*←*bs*_ = 0, *µ*_*bs*_ = 0, 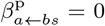 and 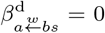, and if group *s* ∈ 𝒮 is at its maximum size then 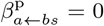 and 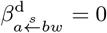 for all *a, b* ∈ 𝒫, *s, w* ∈ 𝒮.

**Figure 1:**
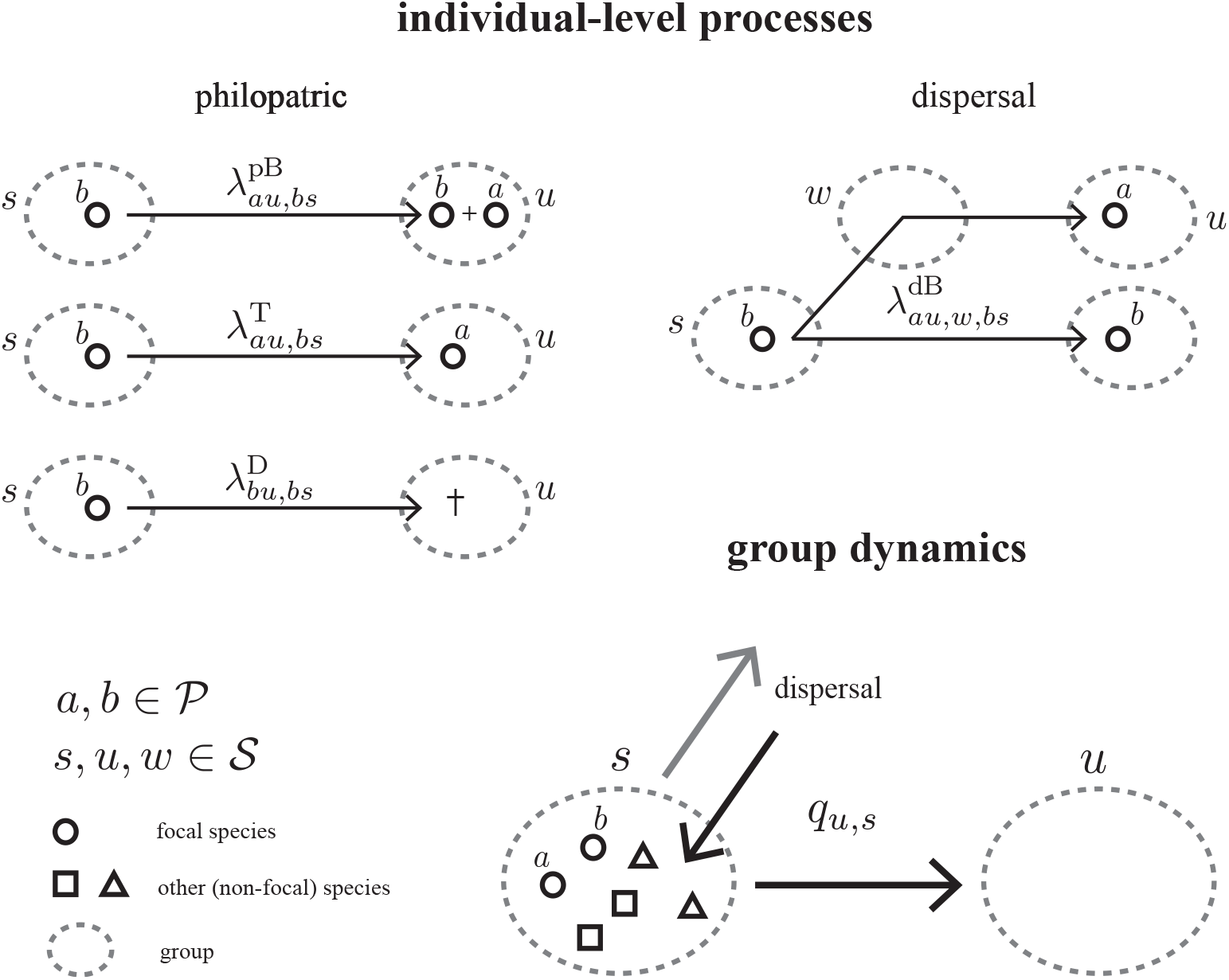
Individual-level processes of the focal species (top panels) and the group dynamics of the meta-community (bottom panels). **Top panel:** As given in Section 3.1, the focal resident species undergoes philopatric individual-level processes of birth, physiological transition and death (top left panel), as well as birth and dispersal (top right panel). The dashed ovals represent groups and circles represent individuals of the focal species. In all processes the parent individual is in physiological state *b* ∈ 𝒫 and inhabits a group in state *s* ∈ 𝒮, and the arrow indicates who was being produced or removed as a result of this process. **Bottom panel:** Group transitions (11)-(14) are caused by the individual-level processes of the focal and all non-focal species as well as fluctuations in external factors. Group transitions determine the change in the composition of groups and the metacommunity at large as given in (10).

Formally, we can express the four basic rates of the focal species as the mappings

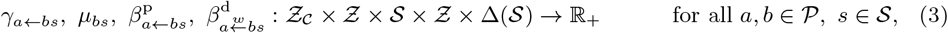

such that, for instance, the first argument in *γ*_*a*←*bs*_(*z*_*bs*_, ***z***, *s*, ***z*, x**) indicates the class-specific phenotype *z*_*bs*_ ∈ *Ƶ*_𝒞_ expressed by the focal individual who undergoes the i-process, and the remaining arguments describe the group of the focal individual and the rest of the metacommunity that affect these i-processes. That is, the second argument ***z*** ∈ *Ƶ* indicates the phenotypic profile of resident individuals of the focal species that inhabit the group of the focal individual and the third argument *s* ∈ 𝒮 gives the state of the entire focal group, namely, the distribution of the focal and non-focal species as well as abiotic factors (Section 2). Likewise, the fourth argument ***z*** ∈ *Ƶ* indicates the phenotypic profile of resident individuals in the focal species outside of the group of the focal individual and the final argument gives the g-state distribution of the entire metacommunity **x** ∈ Δ(𝒮). Moreover, in calculations involving the i-rates (3), it will be useful to use the short-hand notation

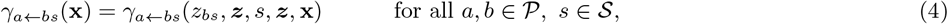

indicating the dependence on the state of the metacommunity. The arguments and the short-hand notation of functions 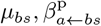 and 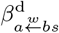, as well as other individual-level functions introduced henceforth, are interpreted identically. Note that the mapping for the total birth rate via dispersal 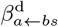 can be obtained by using 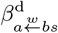.

For ease of presentation and calculation, we will further use a single symbol for all the i-rates (3), and in addition indicate the g-state of the group after the focal individual has undergone the i-process. To this end, and for all *au, bs* ∈ 𝒞, the class-specific physiological transition rate satisfies

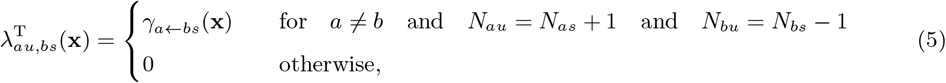

the class-specific death rate satisfies

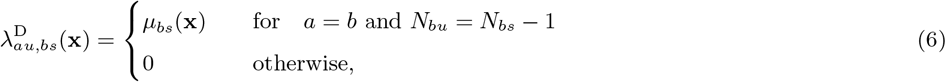

and the class-specific philopatric birth rate satisfies

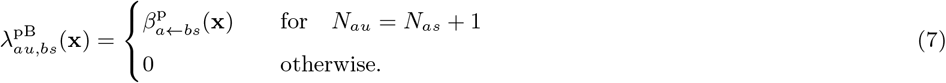

The class-specific conditional birth rate via dispersal satisfies

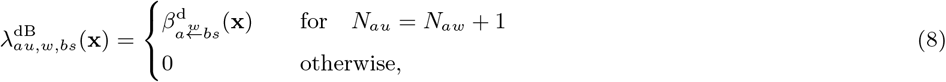

and so the total birth rate via dispersal is

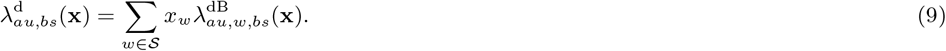

### 3.2 Group state dynamics

The dynamics of the frequency of g-states in the metacommunity **x** ∈ Δ(𝒮) can now be expressed as a system of ODE’s

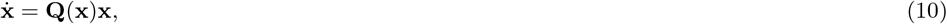

where the dot represents the time derivative and 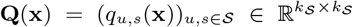 is a group transition-rate matrix where *q*_*u,s*_(**x**) is the rate at which a group in state *s* ∈ 𝒮 transitions to state *u* ∈ 𝒮. Consistency requires that ∑_*u*∈𝒮_ *q*_*u,s*_(**x**) = 0 for all *s* ∈ 𝒮 and hence *q*_*s,s*_(**x**) = − ∑ _*w*≠*s*∈𝒮_ *q*_*w,s*_(**x**) gives the rate at which a group transitions from *s* ∈ 𝒮 to any other g-state (Figure 1). Due to the independence of the i-processes (Section 3.1), we can write

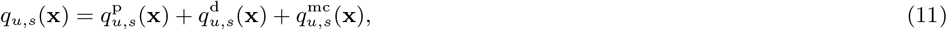

where the group transitions due to philopatric i-processes of the focal species as

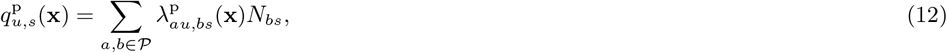

with

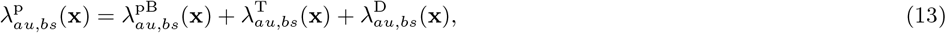

and that due to dispersal as

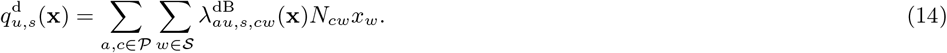

This sums both over all physiological states of the parents and their possible progeny as well, and recall that *N*_*as*_ is a constant specifying the number of individuals *a* ∈ 𝒫 in group *s* ∈ 𝒮 (Section 2). Finally, all other non-focal species processes affecting the metacommunity state dynamics are captured by the rate 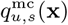.

We note that there are two biologically relevant perspectives on the process generated by (10). First, one can interpret **x** as the frequency distribution of an *infinite collection of groups* and which is taken as the state of the metacommunity (as discussed above). In this view the ODE in (10) describes a deterministic processes on how the state of an infinite collection of groups changes in time. We will call it the deterministic perspective. Second, one can look at a *single group* and interpret the vector **x** = **x**(*t*) as a probability distribution over the state space 𝒮, each element giving the probability of the state the group is in at time *t*. The rates (11) are then interpreted as the rates at which the group transitions from one state in 𝒮 to another state in 𝒮. Taking this perspective, all groups in the metacommunity are assumed to undergo a non-homogeneous continuous time Markov chain (Iosifescu, 2007, chap. 7) with transition matrix **Q**(**x**) that is a function of the state **x** of the metacommunity (hence the non-homogeneity). We call this the probabilistic perspective. In the main text, we follow for ease of presentation the deterministic perspective and for the final results in Section 7 we discuss the probabilistic perspective.

### 3.3 Resident population and reproductive value dynamics

We now describe the population dynamics of the focal species, which needs to be done in the backdrop of the full metacommunity state dynamics (10). Let 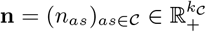 denote the density vector of individuals of the focal species in a monomorphic metacommunity, where *n*_*as*_ = *N*_*as*_*x*_*s*_ defines the density of individuals in class *as* ∈ 𝒞. Because groups are assumed to be well-mixed in the meta-community, the density of individuals refers to the number of individuals per unit space on the scale of groups. The vector 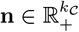 satisfies the system of ODEs

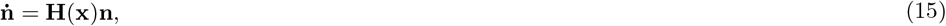

where 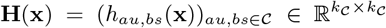 is the resident individual growth-rate matrix. The elements of this matrix are

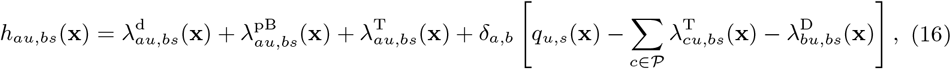

for all *au, bs* ∈ 𝒞, which is the rate at which a focal ‘parent’ individual in class *bs* ∈ 𝒞 produces or removes an ‘offspring’ individual of class *au* ∈ 𝒞 (see for a derivation Appendix A, Section A.1).

Equation (16) gives a representation of the class-specific individual growth-rate, or individual fitness, whose arguments are identically interpreted to those of the i-rates (3)-(4) and it consists of four terms. The first three terms describe the rate at which a parent individual *bs* ∈ 𝒞 produces offspring individuals *au* ∈ 𝒞 via dispersal, philopatric birth and physiological transition as detailed in Section 3.1, and in the case of physiological transition the parent and offspring individuals are interpreted as one and the same individual. The final term in (16) describes the ‘survival’ of the parent individual *bs* ∈ 𝒞 in a changing local environment, where *δ*_*a,b*_ is the Kronecker delta taking value 1 when *a* = *b* ∈ 𝒫 and otherwise 0. More specifically, for *s* ≠ *u* ∈ 𝒮, the final term in (16) gives the rate at which the parent *a* = *b* ∈ 𝒫 survives the group transition from *s* ∈ 𝒮 to *u* ∈ 𝒮, and because *q*_*u,s*_(**x**) contains the i-processes of all individuals including the focal parent, the physiological transition and death of the parent are being subtracted in (16). For *s* = *u* ∈ 𝒮, the survival term reduces to *q*_*ss*_(**x**) = –∑_*w≠s*_ *q*_*w,s*_(**x**) and gives the rate at which the parent *au* = *bs* ∈ 𝒞 is removed from class *au* = *bs* ∈ 𝒞 due to any event that causes the group to transition from *s* = *u* ∈ 𝒮 to any other state in 𝒮, including events caused by the focal individual (and note that in contrast to other processes dispersal may also contribute to fitness for *s* = *u* ∈ 𝒮). The first term in (16) thus contributes to the rate of offspring production through dispersal while the second, third and fourth term together gives the net philopatric fitness. We note that in order to study the dynamics of **n**, the ODE (15) needs to be coupled with the dynamics for the metacommunity state **x** in (10).

The dispersal and net philopatric fitness contributions of individual fitness (16) corresponds to those of previous discrete time models, which are not expressed in terms of individual and group transition rates but directly in terms of the two net fitness contributions (Lehmann et al., 2016, eq. 6, Ohtsuki et al., 2020, eq. 38, and see Appendix A, Section A.1.1). By using (16), we can also represent the individual fitness for models that have been previously considered in the literature. For instance, supposing that the metacommunity is composed only of the focal species that is physiologically unstructured and where groups are identical in terms of abiotic factors, the individual fitness (16) reduces to 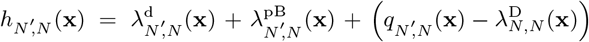 with 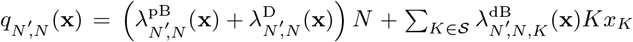 for all *N, N*’ ∈ 𝒮, where the set of g-states 𝒮 = {0, 1, 2, …, *N*_max_} specifies the number of individuals inhabiting a group and where *N*_max_ is the maximum group size (Appendix A, Section A.1.1). Thus, the only processes occurring in each group are death, philopatric birth and birth via dispersal of individuals in the focal species, which is conceptually similar to a number of previous models (Metz and Gyllenberg, 2001; Cadet et al., 2003; Rousset and Ronce, 2004; Lehmann et al., 2006; Alizon and Taylor, 2008; Parvinen, 2013). Further, for well-mixed populations where each ‘group’ is either empty or is inhabited by a single physiologically structured individual of the focal population, the individual fitness reduces to 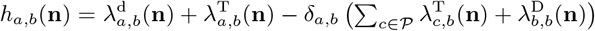, for all *a, b* ∈ 𝒞, where we have 𝒮 = {0} ⋃ 𝒫 and where the population demographic state can be characterized directly in terms of the vector **n** = (*n*_*a*_)_*a*∈𝒫_ of class densities (Appendix A, Section A.1.1). Now, the only processes are physiological transition, death and birth via dispersal because each ‘group’ can contain only a single individual, and which is similar to that of the spatially well-mixed model of Lion (2018a,b, eq. 1) and Priklopil and Lehmann (2020, eq. 3) and conceptual equivalent to standard discrete models of only physiologically structured populations (e.g., Caswell, 2000). This model reduces to an age structured model with *T* discrete age groups by setting 𝒫 = {1, 2, .., *T*} and where newborns are individuals of age group 1 ∈ 𝒫 produced at rate *β*_*b*_ by an individual of age group (class) *b* ∈ 𝒫 so that 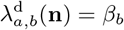 when *a* = 1 and zero otherwise, and where 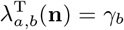 is the constant rate at which an individual in age group *b* ∈ 𝒫 progresses to age group *a* ∈ 𝒫 and where the (average) unit of time an individual spends in an age class *b* ∈ 𝒫 is 1*/γ*_*b*_ (e.g., Li and Brauer, 2008). Setting *γ*_*b*_ = *γ* for all *b* ∈ 𝒫 where age progresses at a steady rate is conceptually similar to the classic discrete-time age-structured models (e.g. Charlesworth, 1994; Caswell, 2000).

Because we are eventually interested in the asymptotic resident species population dynamics, we will utilize the concept of a reproductive value (Fisher, 1930; Taylor, 1990; Rousset, 2004; Grafen, 2006; Lion, 2018b) and use the following population dynamical definition of reproductive values 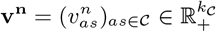

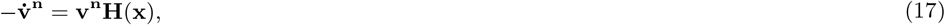

normalized such that **v**^**n**^ · **n** = 1 for all *t*. Because (17) is the adjoint equation (backward in time counterpart) to (15), the superscript in **v**^**n**^ indicates that the reproductive value is an adjoint variable to **n**. Adjoint variables are useful in the analysis of dynamical systems because, by definition, the product of a variable with its adjoint variable is constant in time (e.g. Greiner et al., 1994; Inaba, 2017 in the context of age-structured populations) and one can readily check by applying (15) that **v**^**n**^ · **n** indeed stays constant in time. Similarly to (15), in order to study the dynamics of **v**^**n**^, the ODE (17) needs to be coupled with the dynamics for the metacommunity state **x** in (10). We note that the above definition of reproductive values departs slightly from the standard definition (Taylor, 1990; Lion, 2018b) given as an adjoint to the class frequencies, see further discussion in Section 7.2.

The biological interpretation of **v**^**n**^ is obtained from (17) and is tied to a final value problem where (17) is coupled with a final condition **v**^**n**^(*t*_f_) where *t*_f_ ∈ [*t*, ∞] is some final time: the reproductive value 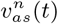 is the probability that a randomly sampled individual at *t*_f_ descends from a single individual of class *as* ∈ 𝒞 at time *t*. Or alternatively, considering an individual of class *as* ∈ 𝒞 at time *t*, 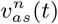 is the *frequency* of individuals at the final time *t*_f_ who are its descendants. In most previous use of reproductive value in evolutionary analysis, as well as in the present paper later on, the final time is usually the distant future (*t*_f_ → ∞) so that 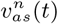 is interpreted as an asymptotic contribution to fitness.

### 3.4 Relatedness dynamics

The remaining variable that appears in (2) and whose dynamics we aim to characterize in the resident population is relatedness. We *define* relatedness between two different individuals *a, b* ∈ 𝒫 inhabiting the same group *s* ∈ 𝒮 at time *t* as the probability *r*_*abs*_ = *r*_*abs*_(*t*) that the two individuals have a common ancestor. In other words, it is the probability that the ancestral lineages of the two individuals coalesce in a common ancestor, and because we assume an infinite number of groups the probability that two individuals sampled from two different groups share a common ancestor is 0. This could be calculated by studying the purely resident i-processes backwards in time (see Michod and Hamilton, 1980 for the original definition and Rousset, 2004 for an analysis of relatedness under a large class of different discrete time scenarios in the island model), but we will here derive it from the forward perspective of the demographic process so as to treat all variables in a similar fashion.

Two individuals that inhabit the same group are related if they either share a parent, who is their common ancestor, or if any of their ancestors shared a parent. Note that in a continuous time Poisson-based model with one birth per unit of time, one of the two individuals that share a parent must be the parent through survival. To find a dynamical equation for relatedness we thus only need to keep track of the ancestors of local parents and to that end we focus on densities of local pairs. We denote a pair of individuals in physiological states *a, b* ∈ 𝒫 that inhabit the same group that is in state *s* ∈ 𝒮 with (*as, bs*) := (*abs*) ∈ ℬ, where ℬ := 𝒫^2^ × 𝒮 is the space of local pairs and whose size is denoted with *k*_ℬ_. Then,

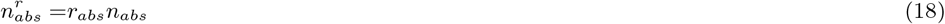

is the density of pairs of individuals who are related and where

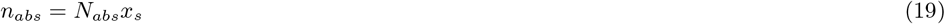

is the total density of pairs *abs* ∈ ℬ in that group. Here,

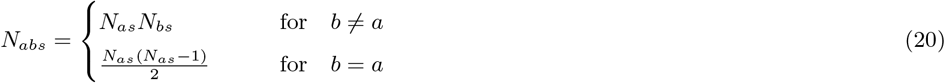

is the number of pairs *abs* ∈ ℬ.

We start by deriving ODEs’ for *n*_*abs*_ and 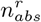, which in turn will allow us to determine *r*_*abs*_. Because we are interested in the ancestry of each pair we will pay special attention to the state and location of ‘parent’ individuals associated to each ‘offspring’. By differentiating (19) we obtain an ODE for the pair dynamics

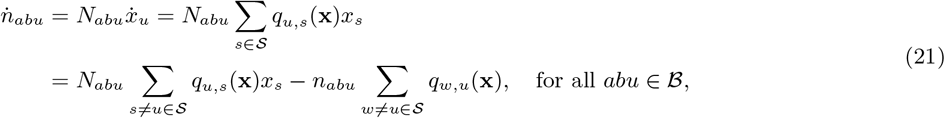

where we used (10), (19) and the fact that ∑_*u*∈𝒮_ *q*_*u,s*_(**x**) = 0 (Section 3.2). By substituting (11)-(14) into (21) and using the relationship *n*_*as*_ = *N*_*as*_*x*_*s*_, we get

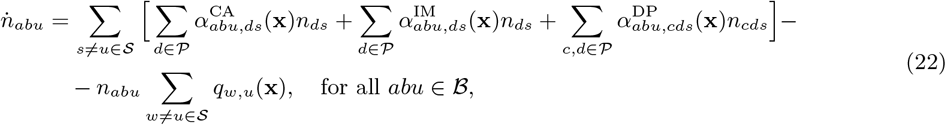

which depends on three rates. First,

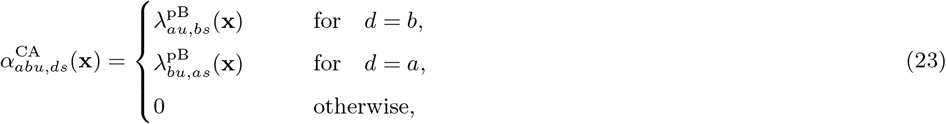

which is the rate at which a single parent individual *ds* ∈ 𝒞 produces offspring pairs *abu* ∈ ℬ by philopatric birth and its own survival and is thus said to be the common ancestor to these pairs (hence the superscript). Second,

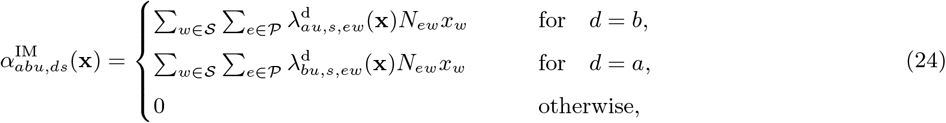

which is the rate at which a single local parent individual *ds* ∈ 𝒞 is being paired up with an immigrant offspring (hence the superscript) to produce an offspring pair *abu* ∈ ℬ. Recall from Section 3.1 that the probability for an immigrant offspring to land in the group of the individual *ds* ∈ 𝒞 depends on the state of the entire group *s* ∈ 𝒮. Finally,

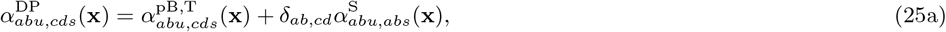

which is the rate at which a local parent pair *cds* ∈ ℬ (two different parents, hence the superscript) produces an offspring pair *abu* ∈ ℬ and this depends on two processes. First,

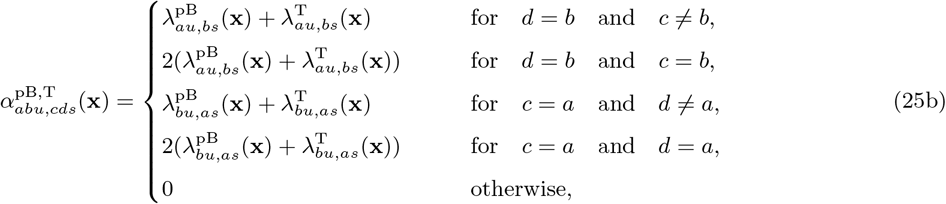

which is the rate at which the local parent pair *cds* ∈ ℬ produces an offspring pair *abu* ∈ ℬ by birth and physiological transition. Because these rates are given for parent pairs, a factor 2 appears whenever both parents can undergo the i-processes that contribute to the birth of the considered offspring pair. Second,

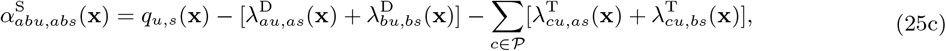

which is the rate at which each existing pair *abs* ∈ ℬ survives upon any i-process occurring in the group (note the similarity of this expression with the ‘survival’ term in (16)). The minus term appears here because parents *a, b* ∈ 𝒫 that die or transition contribute negatively to the formation of offspring pairs *abu* ∈ ℬ. Note that all these rates for production of offspring pairs depend on the rates (5)-(9) at which single individuals undergo demographic i-processes.

The dynamics of 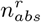 is obtained from (22) by subtracting the immigration term, because immigrants are not related, and by only considering the fraction of local pairs that are related and we get

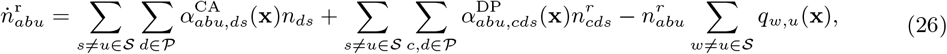

for all pairs *abu* ∈ ℬ. We see from the first two terms in (26) that the density of related pairs *abu* ∈ ℬ increases due to a single local parent producing pairs anew, or, due to two related local parents producing offspring-pairs. From the last term in (26) we see that the density of related pairs *abu* ∈ ℬ decreases due to any event that changes the state of the focal group *u* ∈ 𝒮. The ODE for the vector of relatedness 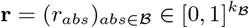 can be given in terms of its elements 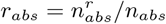 and is obtained by differentiation

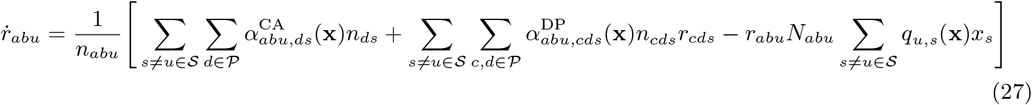

for all *abu* ∈ ℬ, and where we have used (21), (26) and 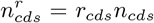. Notice that the rates that appear in (21) and (26) at which pairs *abu* ∈ ℬ are removed due to the group *u* ∈ 𝒮 changing its state, cancel in (27). Hence, the rate at which relatedness changes is the relative rate of producing pairs *abu* ∈ ℬ by a common parent or by parents that share an ancestor, relative to an average rate of producing pairs *abu* ∈ ℬ. We note that similarly to (15) and (17), in order to study the dynamics in (27) it needs to be coupled with the dynamics for the metacommunity state **x** in (10).

By ignoring physiological structure in the focal species, the dynamics in (27) reduces to the expression given in Wild et al. (2009, Supplementary Information, equation S4 and Table S1). Moreover, assuming no class-structure so that all groups consists only of the focal species of constant size *N* and where the vector **x** plays no role, the continuous time dynamical equation for relatedness (27) is simplified further to

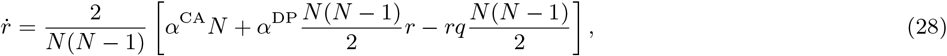

where *q* is the total rate at which events happen in a group. When each birth event is coupled with a death event (i.e., the Moran process, Ewens, 2012), the rates in (28) can be written as

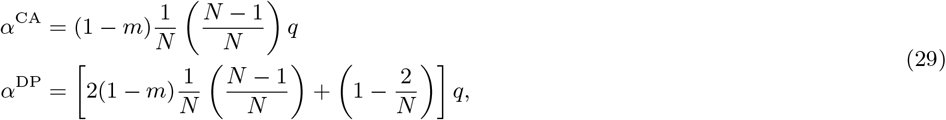

where *m* is the probability that upon a birth event the offspring disperses. Here, *α*^CA^ is interpreted as the rate at which events happen in the group *q* times the probability that the parent individual reproduces locally 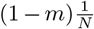 and does not die 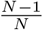. The rate *α*^DP^ is interpreted as the rate at which events happen in the group *q* times the probability that either individual in the pair reproduces locally and does not die 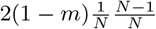, or, the probability that neither individual in the pair dies 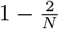.

#### 3.4.1 The jump process for relatedness

Next we represent the continuous time dynamics of relatedness (27) as a discrete time jump-process, with jumps occurring whenever an event changes the state of the group. This representation is useful when only the steady state is considered and also in connecting our result to previous models that have mostly been formulated in discrete time. We thus consider a jump chain (or embedded Markov chain, e.g., Iosifescu, 2007, chapter 8.3) associated to the continuous time Markov chain given in (10) (see further details in Appendix B, Section B.1). An embedded Markov chain is a discrete time Markov chain that records state transitions upon jump times *T*_1_, *T*_2_, …, where each jump time *T*_*k*_ is a random variable at which the focal group undergoes an event either due to local or non-local (via immigration) i-processes. The jump process for relatedness is given in vector form as

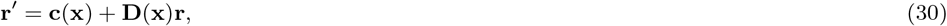

where 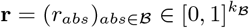 and where **r**′ = **r**(*T*_*k*+ 1_) and **r** = **r**(*T*_*k*_). Here, 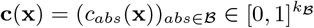 is a vector where entry *c*_*abs*_(**x**) is the probability that a randomly sampled offspring-pair *abs* ∈ ℬ has a common parent in the same group, and hence a common ancestor. This is the coalescence probability and is computed as *c*_*abu*_(**x**) = ∑_*ds*∈𝒞_ *C*_*abu,ds*_(**x**) with

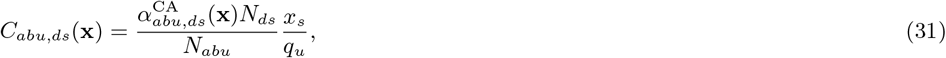

for *s* ≠ *u* ∈ 𝒞 and 0 otherwise, and where *q*_*u*_ = ∑_*w*≠*u*∈𝒮_ *q*_*u,w*_(**x**)*x*_*w*_ is the rate at which groups transition to state *u* ∈ 𝒮. Equation (31) was obtained using the rates (23), and *q*_*u*_ is obtained from the first expression on the right hand side of (21). Equation (31) gives the probability, conditional on the occurrence of a group transition to *u* ∈ 𝒮, that the group was in state *s* ∈ 𝒮 before the transition and a randomly sampled offspring-pair *abu* ∈ ℬ was produced anew by a local parent individual in class *ds* ∈ 𝒞. The matrix 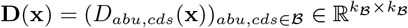 has entries

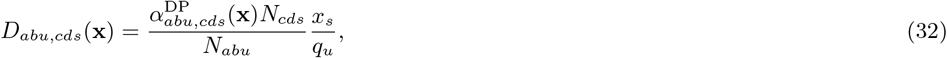

for all *s* ≠ *u* ∈ 𝒞 and 0 otherwise, and gives the probability that, conditional on the occurrence of a group transition to *u* ∈ 𝒮, that the group was in state *s* ∈ 𝒮 before the transition and a randomly sampled offspring-pair *abu* ∈ ℬ was produced by a local parent-pair *cds* ∈ ℬ (and where we used (25)). Note that the probabilities (31)–(32) are both defined to be 0 whenever *N*_*abu*_ = 0.

The recursion (30) along with (31)–(32) reduces to recursion (A.5) of Alizon and Taylor (2008) who considered group size as the only class-structure and hence 𝒞 = 𝒮. The recursion (30) is also conceptually analogous to the recursion for relatedness in the presence of class-structure in the discrete time island model (e.g., Ronce et al., 2000, A.9 for 𝒞 = 𝒫 ×𝒮, Ohtsuki et al., 2020, eq. 33 for 𝒞 = 𝒮). Yet owing to the continuous time nature of our model, expressions (31)–(32) do not exactly match those of the discrete time process since only one individual-level event can occur per unit of time in a continuous time process, while two or more events can occur in discrete time models. Finally, note that in a model with no class-structure and thus no dependence on **x**, the recursion (30) simplifies to

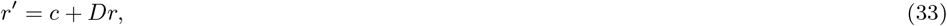

where

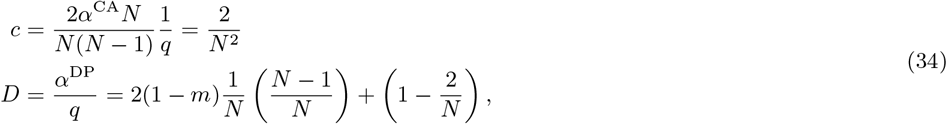

and where the expressions and the interpretation for *α*^CA^, *α*^DP^ and *q* are given in (28)-(29).

### 3.5 Steady states

In the evolutionary analysis that follows we assume that a non-trivial steady state of the metacommunity state dynamics (10) satisfying 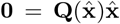 exists, and that it is hyperbolically stable. A steady state is said to be hyperbolically stable if the real part of the dominant eigenvalue of the linearized version of the dynamical system, evaluated at the steady state, is negative and bounded away from zero (Hirsch et al., 1974). This implies that there exists a neighborhood of 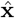 such that for any initial condition **x**(*t*_0_) in this neighborhood where *t*_0_ is some initial time, the vector 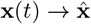 as *t* → ∞.

Because in (15) the population densities of the focal species satisfy *n*_*as*_ = *N*_*as*_*x*_*s*_ for all *as* ∈ 𝒞 where *N*_*as*_ is a constant, then 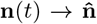 whenever 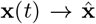 as *t* → ∞. Since 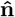 must also satisfy 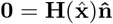, the vector 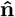 is the dominant right eigenvector of 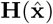. Moreover, because (17) is linear in **v**^**n**^, we have 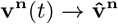 whenever 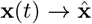 as *t* → ∞ and the steady state for **v**^**n**^ can be found by solving 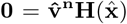. Due to the constraint 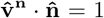, the vector 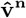 is the (unique) dominant left eigenvector of 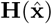. Finally, the steady-state for relatedness **r** can be found by either solving the system of ODEs (27) or the jump-process in (30). By a direct calculation we get

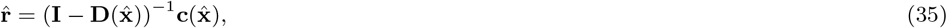

where **I** is the identity matrix and 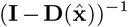 is the inverse matrix of 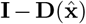 evaluated at 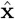 thus allowing to compute relatedness explicitly in terms of resident i-processes (Section 3.1). Because its growth-rate is at most linear in **r** and the vector **c**(**x**) and **D**(**x**) depend only on **x**, we have that 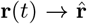 whenever 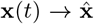 as *t* → ∞. In summary, we have that whenever **x**(*t*) converges to the hyperbolically stable steady state of interest 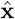 as *t* → ∞, the vectors (**n**(*t*), **v**^**n**^ (*t*), **r**(*t*)) converge to their unique steady state 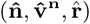 as *t* → ∞.

As a check of the recurrence equations for relatedness, we note that for the homogeneous island model we obtain 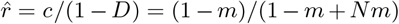 from (34), which recovers, as it should, the relatedness for the discrete time Moran process in the island model (Lehmann et al., 2015, eq. 8-e). Note that at the steady state the relation 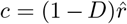 must hold, saying that the probability *c* that two randomly sampled offspring have a common parent must be equal to the probability 1 – *D* that two randomly sampled individuals have two different parents times the probability 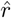 that they are related.

## 4 Mutant-resident dynamics in the metacommunity

We now study the full mutant-resident system and to that end we need to consider that phenotypic expression depends, in addition to class, on the genetic state of the individual. We denote with ***z***_*θ*_ ∈ *Ƶ* the phenotypic profile of a carrier of allele *θ* ∈ {M, R}. Whenever allele *θ* is in an individual in class *as* ∈ 𝒞 the expression of the phenotype is *z*_*θ,as*_ ∈ *Ƶ*_𝒞_ ⊂ ℝ. We thus need to expand the group state space of the metacommunity since the different alleles can be distributed in any way across groups. Following the construction of Section 2, it will be convenient to define the state space of groups for the mutant-resident system as Ω = 𝒮 × ℐ. The set ℐ = {ℐ_*s*_}_*s*∈𝒮_ gives the set of size distributions of mutants of the focal species over all g-states, where each element ℐ_*s*_ ∈ ℐ denotes the set of all possible size distributions in a group *s* ∈ 𝒮. This is defined as ℐ_*s*_ = {ℐ_*as*_}_*a*∈𝒫_ with 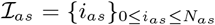 where *i*_*as*_ is the number of mutants in physiological state *a* ∈ 𝒫 that inhabit a group *s* ∈ 𝒮. We note that an element *i* ∈ ℐ_*s*_ is a *vector* giving the number of mutants in each physiological state in a group *s* ∈ 𝒮 (the set ℐ_*s*_ has potentially many elements because a group *s* ∈ 𝒮 can be in different genetic states). The number of residents in a group can be obtained simply by subtracting the number of mutants from the total number of individuals e.g. *N*_*as*_ – *i*_*as*_ is the number of residents *a* ∈ 𝒫 in a group *s* ∈ 𝒮. We note that while 𝒮 gives the demographic structure of the entire group including non-focal species and abiotic factors, the set ℐ refers only to the genetic structure of the focal species.

Similarly to Section 3, we need to track the dynamics of mutant-resident group state frequencies in order to characterize the mutant-resident dynamics. We represent the mutant-resident state of the metacommunity with a vector **y** = (*y*_*si*_)_*si*∈Ω_ ∈ Δ(Ω) of length *k*_Ω_, where *y*_*si*_ is the frequency of groups in mutant-resident state (*s, i*) := *si* ∈ Ω and hence ∑_*si*∈Ω_ *y*_*si*_ = 1 and 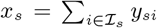 for all *s* ∈ 𝒮 where **x** = (*x*_*s*_)_*s*∈𝒮_ ∈ Δ(𝒮). All the upcoming rates describing the mutant-resident metacommunity dynamics are identical to those given for the resident dynamics (Section 3), the only difference is that it takes into account the genetic structure of the group and the metacommunity at large.

### 4.1 Individual-level processes

The rates at which i-processes of the focal species in the mutant-resident metacommunity occur are defined, similarly to the resident metacommunity in Section 3.1, as mappings

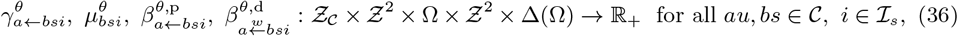

where *θ* ∈ {M, R} specifies whether it is a mutant or a resident who undergoes the process. We also use a shorthand notation 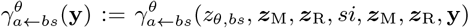 where the arguments are interpreted similarly to Section 3.1, and the interpretation and shorthand notation for the other rates in (36) is similar. Moreover, we assume that dispersed offspring land into a group *wi* ∈ Ω with a probability *x*_*w*_, that is, independent of its genetic state, and hence we will write 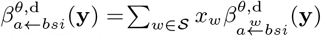. The constrains for the i-rates (36) are also analogous to Section 3.1. Finally, and for all *au, bs* ∈ 𝒞 and *j* ∈ ℐ_*u*_, *i* ∈ ℐ_*s*_, the class-specific physiological transition rate satisfies

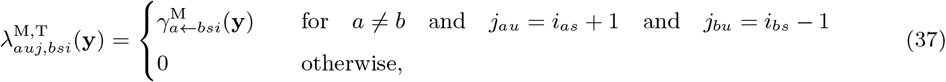

the class-specific death rate satisfies

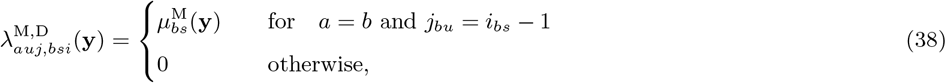

and the class-specific philopatric birth rate satisfies

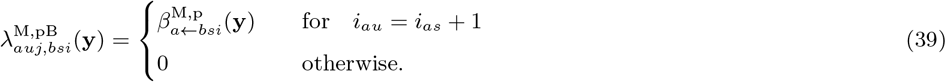

The class-specific conditional birth rate via dispersal satisfies

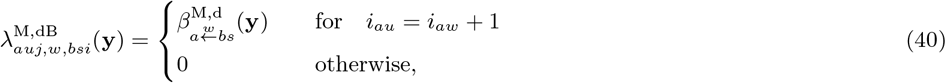

and so the total birth rate via dispersal is

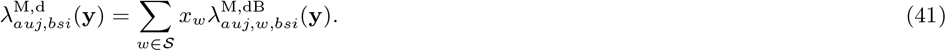

The rates for resident individuals are defined similarly, one only needs to replace the number of mutants *i*_*as*_ with the number of resident *N*_*as*_ – *i*_*as*_ for all *as* ∈ 𝒞.

### 4.2 Group state dynamics

The mutant-resident metacommunity dynamics is expressed analogously to the resident metacommunity dynamics (10) as a system of ODEs’

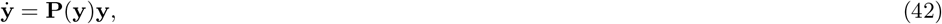

where 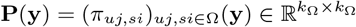 is the group transition-rate matrix with

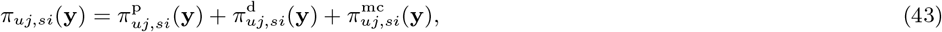

giving the rate at which a group in state *si* ∈ Ω transitions to state *uj* ∈ Ω, and where ∑_*uj*∈Ω_ *π*_*uj,si*_ = 0 for all *si* ∈ Ω. Here, 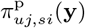 are the philopatric and 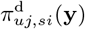 the dispersal group transition rates of the focal species, and where all other transition rates due to other species and abiotic factors are given in 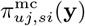. The transition rates due to the focal species can be expressed in terms of i-processes given in (37)-(41) by summing over all physiological states of the parents and their possible progeny, that is, we have

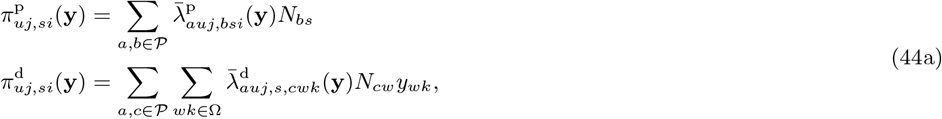

for all *au, bs* ∈ 𝒞, *j* ∈ ℐ_*u*_, *i* ∈ ℐ_*s*_ and *θ* ∈ {M, R}, where

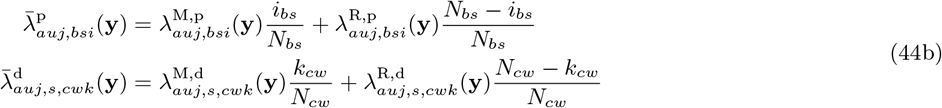

with

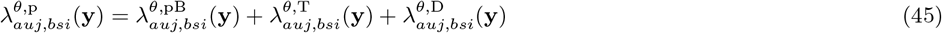

are the rates of an average individual where the average is taken over the allelic states M and R.

### 4.3 Mutant-resident population dynamics

Consider the focal species in the metacommunity and let 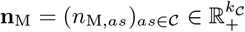 denote the mutant density vector where 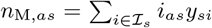 is the density of mutant individuals in class *as* ∈ 𝒞, and let 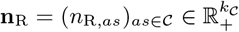 denote the resident density vector where 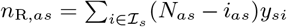 is the density of resident individuals in class *as* ∈ 𝒞. The mutant-resident density dynamics is given by

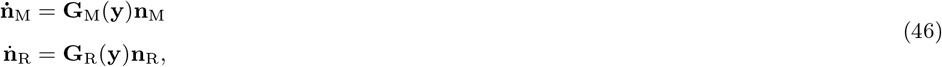

where 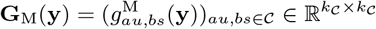 is the mutant fitness matrix with elements

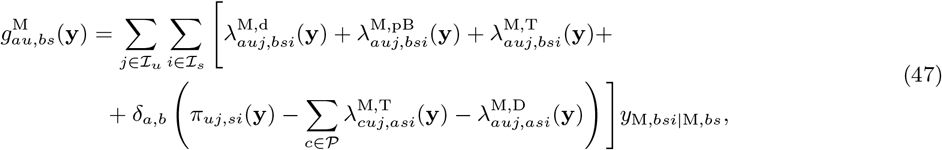

giving the rate at which a single mutant in class *bs* ∈ 𝒞 produces mutants in class *au* ∈ 𝒞, and where 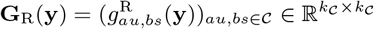 is the resident fitness matrix with elements

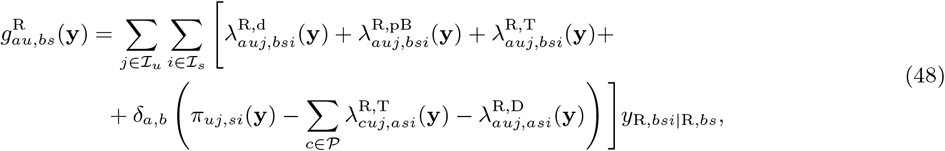

giving the rate at which a single resident in class *bs* ∈ 𝒞 produces residents in class *au* ∈ 𝒞. Here, *y*_M,*bsi*|M,*bs*_ = (*i*_*bs*_*y*_*si*_)*/n*_M,*bs*_ denotes the probability that a mutant *bs* ∈ 𝒞 inhabits a group in genetic state *i* ∈ ℐ_*s*_ and *y*_R,*bsi*|R,*bs*_ = ((*N*_*bs*_ – *i*_*bs*_)*y*_*si*_)*/n*_R,*bs*_ denotes the probability that a resident *bs* ∈ 𝒞 inhabits a group in genetic state *i* ∈ ℐ_*s*_ (Appendix A, Section A.2.1). Note that similarly to the resident species population dynamics, all the densities of all species are functions of the mutant-resident metacommunity state **y** and hence in order to study the dynamics of **n**_M_ and/or **n**_R_ in (46) they need to be coupled with the dynamics of **y** in (42).

### 4.4 Relative mutant-resident population dynamics

To analyse the spread of the mutant allele, it is convenient to rewrite the mutant-resident population dynamics (46)-(48) by using two new vectors. First, with a slight abuse of notation, we use a vector 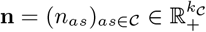 of total densities where *n*_*as*_ = *N*_*as*_*x*_*s*_ for all *as* ∈ 𝒞 is the total density of mutant and resident individuals in class *as* ∈ 𝒞. Second, we use a vector 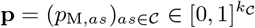 to describe the within-class mutant frequencies where *p*_M,*as*_ = *n*_M,*as*_*/n*_*as*_ for all *as* ∈ 𝒞 is the proportion (frequency) of mutants within class *as* ∈ 𝒞. Note that the pair of vectors (**n**_M_, **n**_R_) can be expressed in terms of (**n, p**), and *vice versa*, because (*n*_M,*as*_, *n*_R,*as*_) = (*p*_M,*as*_*n*_*as*_, (1 – *p*_M,*as*_)*n*_*as*_) for all *as* ∈ 𝒞 and (*n*_*as*_, *p*_M,*as*_) = (*n*_M,*as*_ + *n*_R,*as*_, *n*_M,*as*_*/*(*n*_M,*as*_ + *n*_R,*as*_)) for all *as* ∈ 𝒞. The latter representation is common in population genetics (e.g., Nagylaki, 1992).

The vector of density dynamics satisfies

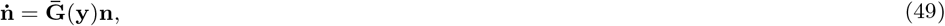

where 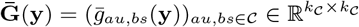 is the average mutant-resident fitness matrix where each element

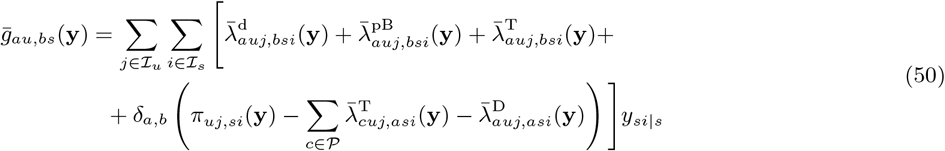

gives the rate at which an average individual in class *bs* ∈ 𝒞 produces individuals of class *au* ∈ 𝒞. The average is taken over the allelic states M, R and over all the genetic states the group it inhabits can be in, so that *y*_*si*|*s*_ = *y*_*si*_*/x*_*s*_ is the probability that the group *s* ∈ 𝒞 that the average individual inhabits is in genetic state *i* ∈ ℐ_*s*_ (see Appendix A, Section A.2.2). The vector of within-class mutant frequency dynamics satisfies

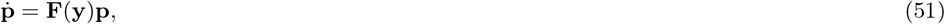

where

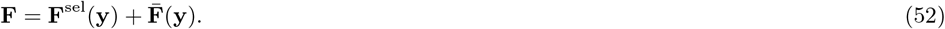

Here, 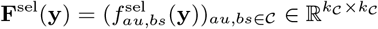 and 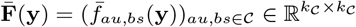 have elements, respectively, given by

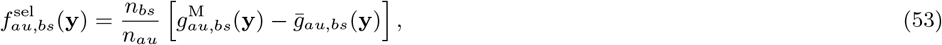

and

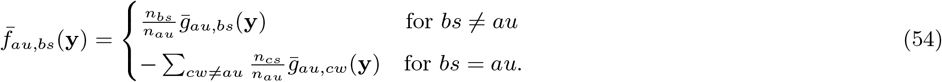

Because the rate in (53) describes differential growth due to different individuals having different phenotype it captures the effect of selection on allele frequency change (hence the superscript). And because the rate in (54) depends only on the average phenotype it captures effects on allele frequency change due to individuals in different classes having different average growth rates. We have added a bar analogously to (49) as it depends only on the average phenotype where the average is taken over the allelic states (see further discussion in Section 8). A similar partitioning of **F** was given for well-mixed populations in Lion (2018b, Appendix 3), and Priklopil and Lehmann (2020, eq. 8a) and the changes due to non-selective forces have been referred to as changes due to “transmission” in population genetics (Kirkpatrick et al., 2002). We note that because **n** and **p** are functions of the state of the mutant-resident metacommunity **y**, in order to study the relative mutant-resident population dynamics of **n** and/or **p**, (49) and/or (54) need to be coupled with the dynamics of **y** in (42).

### 4.5 Average mutant frequency

The total proportion (frequency) of mutants in the metacommunity given as an arithmetic mean is defined as

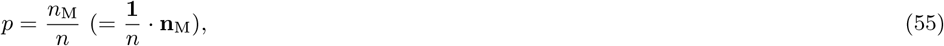

where *n*_M_ = ∑_*as*∈𝒮_ *n*_M,*as*_ is the total mutant density, *n* = ∑_*as*∈𝒮_ *n*_*as*_ is the total mutant and resident density in the focal species (with a slight abuse of notation) and **1** is a vector of all 1 of length *k*_𝒞_. The dynamics of *p* in (55) is obtained by differentiation and by using Sections 4.2-4.4, and it can be written as

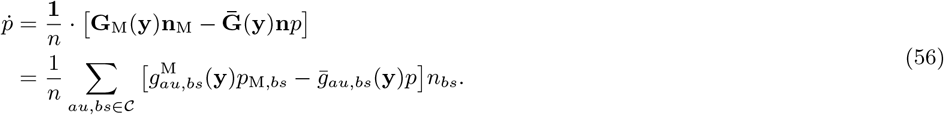

In order to study its dynamics it must be coupled with the dynamics of **y** in (42).

## 5 Mutant-resident dynamics for similar phenotypes

We now study mutant-resident dynamics for closely similar phenotypes, with a particular focus on the change in the arithmetic mean allele frequency (55). To that end, we set ***z***_M_ = ***z***_R_ + *δ****η*** where the vector 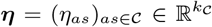 gives the direction of the deviation (where ***z***_R_ = ***z*** ∈ *Ƶ*) and *δ* is a small parameter. To evaluate the mutant-resident dynamics for small *δ*, we Taylor expand about *δ* = 0 the dynamical equations for **y, n** and **p** in (42), (49) and (51), respectively. To facilitate the Taylor expansion, we use the following consistency relation relating mutant-resident i-processes (Section 3.1) to the resident i-processes (Section 4.1):

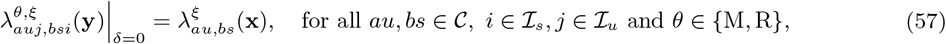

and for ξ ∈ {d, pB, T, D}, and we will also use 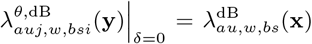. These consistency relations say that whenever mutants and residents express identical phenotypes, the i-processes are independent of the genetic structure of the metacommunity. From (57), it directly follows that

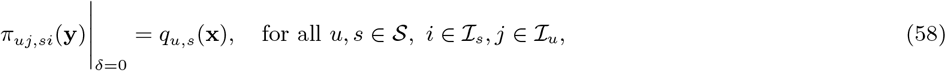

and

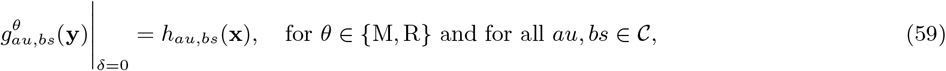

and consequently the growth-rates in (49) and (51) satisfy

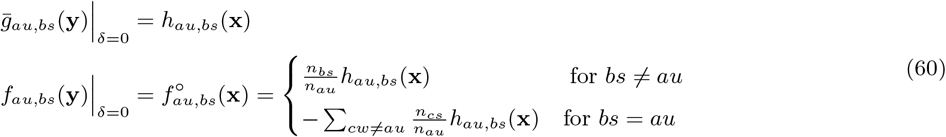

for all *au, bs* ∈ 𝒞.

It follows from the above relations that the dynamics for **y, n** and **p** for small *δ* can be written as

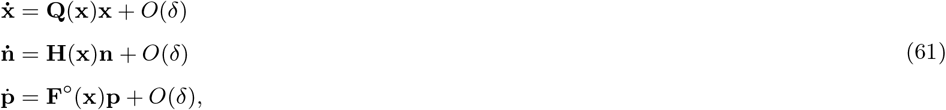

where we used (58) and (60) and where 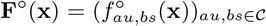 is the relative fitness matrix for mutants (when *δ* = 0) with elements as defined in (60). Note that because the dynamics of **n** and **p** depend only on **x** up to order *O*(1), we have represented the dynamics of the genetically explicit vector **y** by using the genetically implicit vector **x** whose dynamics in (61) up to order *O*(1) follows from (42), (58) and 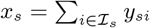 (the dependency on **y** in (61) comes only via the terms of order *O*(*δ*)). The dynamics of the arithmetic mean mutant frequency *p* in (55) for small *δ* is obtained by Taylor expansion and by using (59)-(60), in which case (56) reduces to

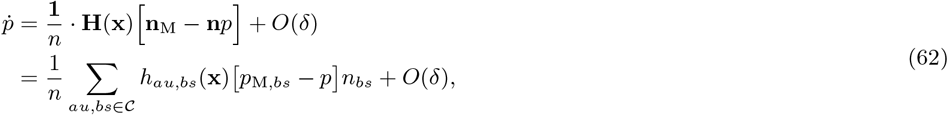

which must be coupled with the dynamics for **p** and **x** in (61) and where the dependency on **y** comes only via *O*(*δ*).

The remaining aim of this paper is to prove the “invasion implies substitution”-principle by way of applying geometric singular perturbation theory developed for *fast-slow ODE systems* in Fenichel (1979). A fast-slow ODE system in a standard form (Fenichel, 1979; Kuehn, 2015; Wechselberger, 2020), which is relatively easy to analyze, is a system of ODEs where a model parameter tunes the rate at which the various dynamical variables operate: for small parameter values *δ* some variables are fast variables in that their Taylor expansion about *δ* = 0 is dominated by *O*(1), while others are slow variables and their Taylor expansion about *δ* = 0 is dominated by *O*(*δ*). We see from (61), however, that for small *δ* the dynamics of the vectors **x, n** and **p**, as well as the arithmetic mean mutant frequency *p* in (62), are all dominated by the terms *O*(1) thus all having a phase of fast dynamics. As a consequence, the mutant-resident system is not readily in the desired (standard) fast-slow form.

In order to construct an appropriate fast-slow system, we proceed in three steps. First, we study the purely fast dynamics of the within-class mutant frequency vector **p** where *δ* = 0, and find its steady state 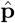 (Section 5.1). Second, by using the properties of 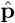 we find a slow variable that can be used, on a suitably chosen slow timescale, as a proxy for the arithmetic mean mutant frequency (Section 5.2). Finally, we study the slow dynamics of this slow variable and calculate all the necessary fast variables (Section 5.3) that define the complete mutant-resident fast-slow system which will be then studied in Section 6.

### 5.1 Fast dynamics of the within-class mutant frequency

To study the fast dynamics of the vector **p** we set *δ* = 0 in the third equation in (61), which gives

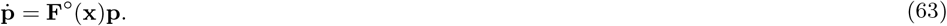

Because the rows of **F°** sum to 0 it is well-known that the steady state satisfies

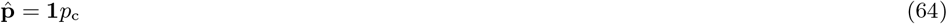

for some constant *p*_c_ ∈ ℝ (e.g., Lessard and Soares, 2018). This implies that the within-class mutant frequencies at the steady state are all equal 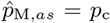 for all *as* ∈ 𝒞. For biologically meaningful values we need *p*_c_ ∈ [0, 1], and the exact value of *p*_c_ that *p*_M,*as*_(*t*) for all *as* ∈ 𝒞 converges to as *t* → ∞ depends on the initial condition (**x**(*t*_0_), **p**(*t*_0_)) for some initial time *t*_0_. The system (63) thus contains infinite number of steady states (equilibria) and the biologically meaningful values lie on a line [0, 1]. This has two important consequences. First, any linear combination of within-class frequencies, at the steady state (64), is equal to *p*_c_. More specifically, every arbitrarily weighted average mutant frequency

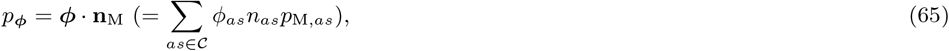

where ***ϕ*** = (*ϕ*_*as*_)_*as*∈𝒞_ is some arbitrary vector of weights normalized such that ***ϕ*** · **n** = 1, must satisfy

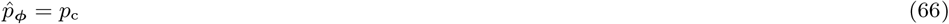

at the steady state where 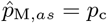 for all *as* ∈ 𝒞. This is in particular true also for the arithmetic mean frequency *p* in (55) where ***ϕ*** = **1***/n*. This implies that every average mutant frequency *p*_***ϕ***_(*t*) → *p*_c_ as *t* → ∞, and where the exact value of *p*_c_ depends on the initial condition (**x**(*t*_0_), **p**(*t*_0_)). Second, the *k*_𝒞_ -dimensional vector **p** is, at the steady state (64), a 1-dimensional vector 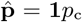 because it can be spanned by using a single vector **1***p*_c_ (Hirsch et al., 1974). This will play an important role in the proof of the “invasion implies substitution”-principle (Section 7). Finally, we want to note that so far we have analyzed the fast dynamics of **p** and *p*_***ϕ***_ dominated by *O*(1) only; in later sections we will see that these variables have also a phase of slow dynamics and this will be analyzed in later sections (see in particular Section 6).

### 5.2 Reproductive value weighted mutant frequency

In the previous Section 5.1 we found that for *δ* = 0 and for any weights ***ϕ***, the average mutant frequency (65) satisfies *p*_***ϕ***_(*t*) → *p*_c_ as *t* → ∞ with the exact value of *p*_c_ depending on the initial condition (**x**(*t*_0_), **p**(*t*_0_)). If one were then to find weights ***ϕ*** for which *p*_***ϕ***_(*t*) = *p*_c_ for all *t* ≥ *t*_0_, then such an average frequency would stay constant in the fast dynamics where *δ* = 0, and hence would be a slow variable for a system where *δ* is small but non-zero (a pre-requisite to construct a standard fast-slow system). We can achieve this by setting ***ϕ*** = **v**^**n**^ where the weights **v**^**n**^ for non-zero *δ* are (with a slight abuse of notation) defined as average reproductive values, where the average is taken over allelic states (Priklopil and Lehmann, 2020, see also Lion, 2018a,b), and is thus the mutant-resident reproductive value analogue to the one defined for the resident population in Section 3. Such average individual reproductive values satisfy

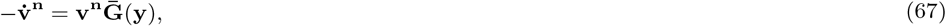

for any *δ*, with the normalization **v**^**n**^ · **n** = 1 where **n** is given by (49). By defining a reproductive values weighted mutant frequency (see (65)) as

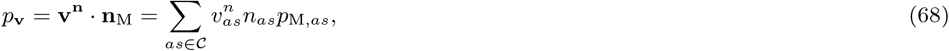

its dynamics is obtained by differentiation and satisfies

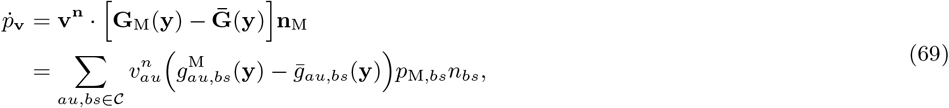

for any *δ*. Indeed, because 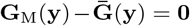 for *δ* = 0 (59)-(60), the Taylor expansion of (69) about *δ* = 0 is dominated by *O*(*δ*) and hence the reproductive value weighted average mutant frequency *p*_**v**_ is a slow variable. We note that since (69) is valid for arbitrary *δ* and thus arbitrary strength of selection, it must be coupled with the dynamics of **y** in (42).

### 5.3 Slow dynamics of the weighted mutant frequency

We here study the slow dynamics of the weighted mutant frequency *p*_**v**_ in (68)-(69), with a specific aim to identify all the dynamical variables that it depends on in order to construct a complete fast-slow mutant-resident system. To that end, we Taylor expand (69) about *δ* = 0 and then change the timescale to slow time while letting *δ* go to 0. To this end, we introduce a slow time variable *τ*, set *τ* = *δt* from which we get the relation 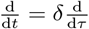, and by letting *δ* go to 0 we get

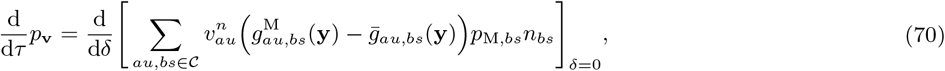

giving the rate at which the reproductive valued weighted frequency changes in a pure slow time *τ* where *δ* = 0. Note that in the fast timescale *t*, the above expression is nothing else than the *O*(*δ*)-term of the Taylor expansion.

Next, we take the derivative in (70) in such a way that it allows us to arrive at an expression for the directional selection coefficient as given in (2). To do this, we must take separate (partial) derivatives with respect to phenotypes of individuals according to the ‘spatial’ relationship between the focal individual and other individuals from the focal species who affect its fitness either via interactions or directly (the effect on itself). For ease of presentation, we label the phenotypes and write the individual fitness (16) as

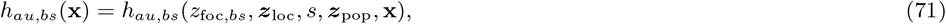

where *z*_foc,*bs*_ (and ***z***_foc_) indicates the phenotype of the focal individual (hence the subscript), ***z***_loc_ indicates the phenotype of the local group members of the focal individual (hence the subscript) but excluding the focal individual, and ***z***_pop_ indicates the phenotype of individuals outside of the group of the focal individual (i.e. population/species at large, hence the subscript). We emphasise that all individuals in the resident metacommunity express the resident phenotype ***z*** ∈ *Ƶ* and that these are just labels to distinguish the spatial relationship of individuals to the focal individual according to which the different partial derivatives will be taken in Section 5.3.1. Likewise, we write *g*_*au,bs*_(**y**) = *g*_*au,bs*_(*z*_M,*bs*_, ***z***_M,loc_, ***z***_R,loc_, *si*, ***z***_M,pop_, ***z***_R,pop_, **y**) for a focal mutant and 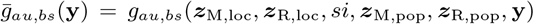 for an average individual where the notation and interpretation of 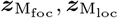 and 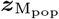 (and 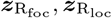 and 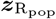) for the mutant-resident metacommunity is similar.

#### 5.3.1 Properties of fitness

To take the derivative in (70) we apply a property that relates, for small *δ*, mutant-resident fitnesses (47)-(48) to the resident fitness (16). This property follows from the so-called generalized law of mass action in situations where weak selection results from small differences in phenotype (Diekmann et al., 2001; Meszéna et al., 2005; Dercole, 2016, for discussion on the different forms of weak selection see Wild and Traulsen, 2007; Lehmann and Rousset, 2014). The generalized law of mass action says that the first-order perturbation of the mutant phenotype of all mutant individuals in a spatially well-mixed mutant-resident species, is equal to the first-order perturbation of the phenotype of all individuals in a purely resident species when multiplied by the frequency of mutant individuals. This is computationally convenient because instead of considering full distributions of individuals only population means need to be calculated. This property holds for non-linear environmental feedbacks and is secured by assuming that all individuals undergo pairwise interactions in continuous time and that individuals with the same phenotype are exchangeable (Dercole, 2016). Below we will provide an analogue of this property for the present metacommunity model.

For small but non-zero *δ*, the mutant fitness matrix **G**_M_(**y**) is related to the resident fitness matrix **H**(**x**) in the following way. First,

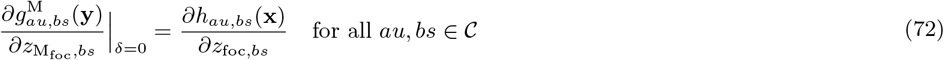

and where all other partial derivatives with respect to 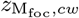 for *cw* ≠ *bs* ∈ 𝒞 are 0. Second,

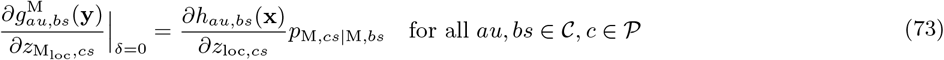

where all other partial derivatives with respect to 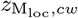 for *w* ≠ *s* ∈ 𝒮 are 0, and finally,

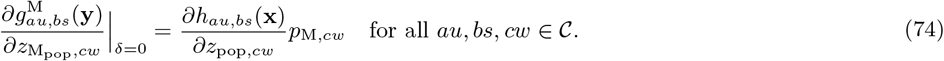

In (73) we introduced the *conditional* within-group mutant frequency 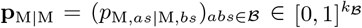 defined as

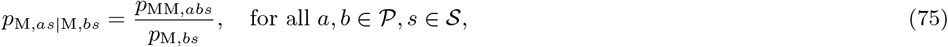

where *p*_MM,*abs*_ is an element of the vector of (within-group) mutant-pair frequencies 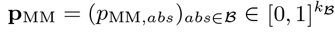 defined as

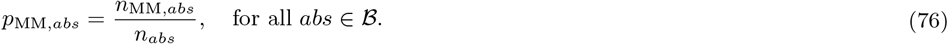

Here, *n*_MM,*abs*_ = ∑_*i*∈I_ *i*_*as*_*i*_*bs*_*y*_*si*_ denotes the density of mutant-pairs *abs* ∈ ℬ and *n*_*abs*_ is the total density (with a slight abuse of notation) of pairs *abs* ∈ ℬ, that is, the total density of pairs independent of their phenotype. The conditional mutant frequencies **p**_M|M_ thus give the frequency of mutants conditional that another (randomly sampled) individual in the group is a mutant.

In (72), the partial derivative is taken with respect to the focal individual itself and because any mutant individual is (trivially) a mutant with probability 1, the proportionality coefficient is simply 1. In other words, the ‘frequency’ of a mutant allele in a single mutant individual is 1. The derivative on the right-hand-side measures (when multiplied by *δ*) the additional offspring the focal individual produces per unit of time due to the focal individual itself expressing the mutation (‘direct fitness effect’). In (73), the partial derivative with respect to all local individuals is proportional to the conditional within-group mutant frequency in class *cs* ∈ 𝒞, *p*_M,*cs*|M,*bs*_, conditional on the focal individual being a mutant *bs* ∈ 𝒞. This property reflects the fact that genetic correlations between conspecifics build up within groups of finite size, and the derivative on the right-hand-side measures (when multiplied by *δ*) the additional offspring the focal individual produces per unit of time due to interactions with all local individuals that express the mutation (‘indirect fitness effect’). Finally, the partial derivative with respect to non-local individuals in (74) is proportional to the (unconditional) within-class mutant frequency because no genetic correlations build up between members in different groups in the infinite island model (see, e.g., Rousset, 2004 for the finite island model).

Using (72)-(74) and analogues properties for (mutant-resident) resident fitness (not shown here), the average fitness matrix 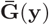 is related to the (pure resident) resident fitness matrix **H**(**x**) as

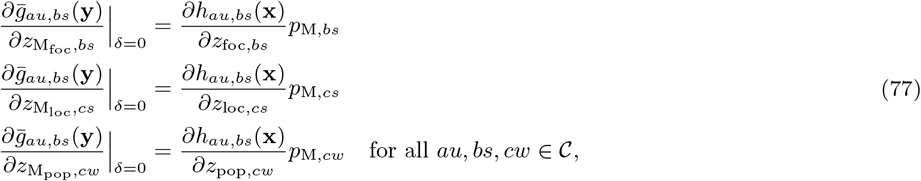

and where all other partial derivatives with respect to 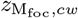 for *cw* ≠ *bs* ∈ 𝒞 and with respect to 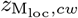 for *w* ≠ *s* ∈ 𝒮 are all 0. Notice that all proportionality coefficients are (unconditional) within-class mutant frequencies. In the first relation in (77) this is because an average individual is a mutant with probability given by the within-class mutant frequency, in the second relation this is because no genetic correlations arise for an average individual and in the final relation this is because no genetic correlations arise between non-group members.

#### 5.3.2 Slow dynamics of the weighted mutant frequency and the fast variables

Taking the derivative in (70) and partitioning it according to the different individuals as discussed above, we obtain

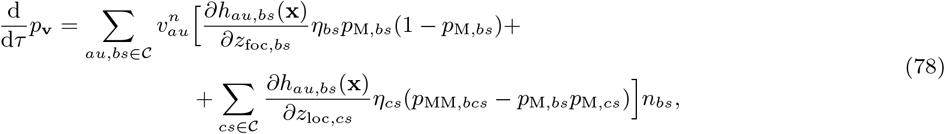

where we used the definition for directional derivatives, the properties (72)-(74), (77) and (75). We can see from (78) that the dynamics of the slow weighted mutant frequency *p*_**v**_ depends on the dynamics of **x, n, p, v**^**n**^ and **p**_MM_. This is convenient because instead of analyzing the dynamics of the large system for **y** it is enough to study the simpler dynamics of **x, n, p, v**^**n**^ and **p**_MM_ (and as will be shown below). Because we have already derived dynamical equations for **x, n** and **p**, and in (61) showed that they are dominated by *O*(1) and so is **v**^**n**^ because under the consistency relation (59) its dynamics reduces to (17), the next task is to find the equation for a vector of mutant-pair frequencies **p**_MM_ (Section 5.3.3).

#### 5.3.3 Mutant-pair frequencies

In order to see whether **p**_MM_ defined in (76) is a fast or slow variable, we study its dynamical equation for *δ* = 0 (Appendix B, Section B.2). The dynamics of **p**_MM_ = (*p*_MM,*abs*_)_*abu*∈ℬ_ is given as

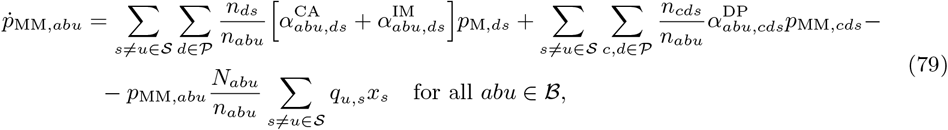

which can be written in vector form as

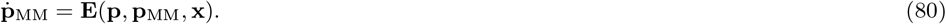

Here, 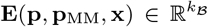 is a vector of rates at which **p**_MM_ changes, and because the elements are non-zero the vector of mutant-pair frequencies **p**_MM_ is dominated by terms of order *O*(1) and is hence a fast variable (it is to be seen whether it changes also in slow time, Section 6).

To characterise for *δ* = 0 the steady state of **p**_MM_ we focus for convenience on the equivalent discrete time jump-process dynamics, which is given by

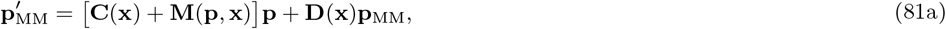

where **C**(**x**) and **D**(**x**) are as in (30), and 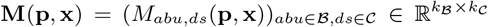 where *M*_*abu,ds*_(**p, x**) = ∑_*ew*∈𝒞_ *M*_*abu,ds,ew*_ (**x**)*p*_M,*ew*_ and

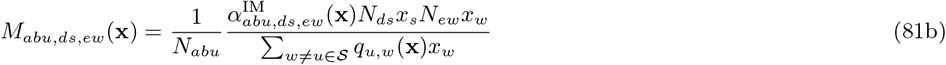

gives the conditional probability that a randomly sampled mutant-pair *abu* ∈ ℬ was produced when a local mutant parent *bs* ∈ 𝒞 paired up with a new immigrant mutant offspring *ew* ∈ 𝒞 (note the dependency on the mutant frequency in (81)). For all pairs *abu* ∈ ℬ we have

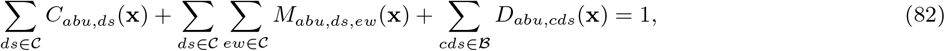

saying that with probability 1 each pair of individuals has either a common parent, different local parents, or non-local parents (the rows of the matrices **C**(**x**), **D**(**x**) and **M**(**p, x**) sum to 1).

The steady state solution 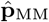 for *δ* = 0 is obtained by solving 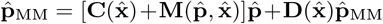 in (81) evaluated at 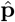 and 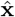, and we get

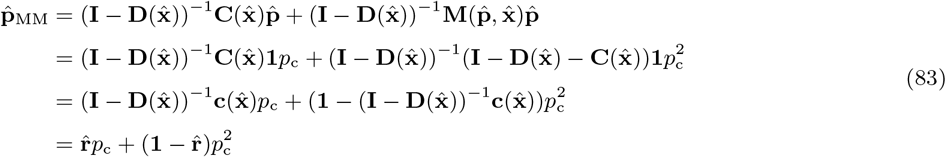

where we used (64), (82) and (35), respectively, and where *p*_c_ ∈ [0, 1] is a constant that depends on the initial condition as discussed in Section 5.1 (we can also obtain this solution by solving (79)-(80)). We have thus obtained that whenever 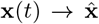 as *t* → ∞ (Section 3.5), then 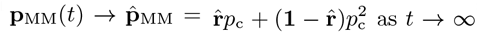 as *t* → ∞. Moreover, 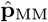 can be expressed in terms of the steady state value for relatedness 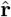 (Section 3.5) and *p*_c_ (Section 5.1), and hence its general solution consist of infinite (curve of) steady states (equilibria) because it is a quadratic function of *p*_c_ ∈ [0, 1]. Finally, we note that we have made no assumptions on the relationship between within-group mutant pair frequencies **p**_MM_(*t*), relatedness **r**(*t*) and the average mutant frequency *p*_***ϕ***_(*t*) for *t* < ∞: their relationship given in (83) is obtained only as a limit when *t* → ∞.

## 6 Mutant-resident dynamics as a fast-slow system

We are now ready to analyse the fast-slow ODE system where mutant and resident individuals have closely similar phenotypes (*δ* is small). Owing to Section 5, the complete fast-slow system for the mutant-resident dynamics is given in fast time *t* as

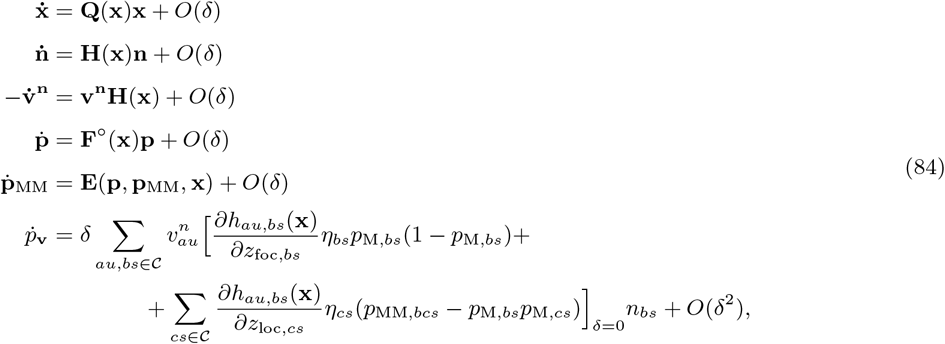

which, on using d*/* d*t* = *δ* d*/* d*τ* can be equivalently written in slow time *τ* = *δt* as

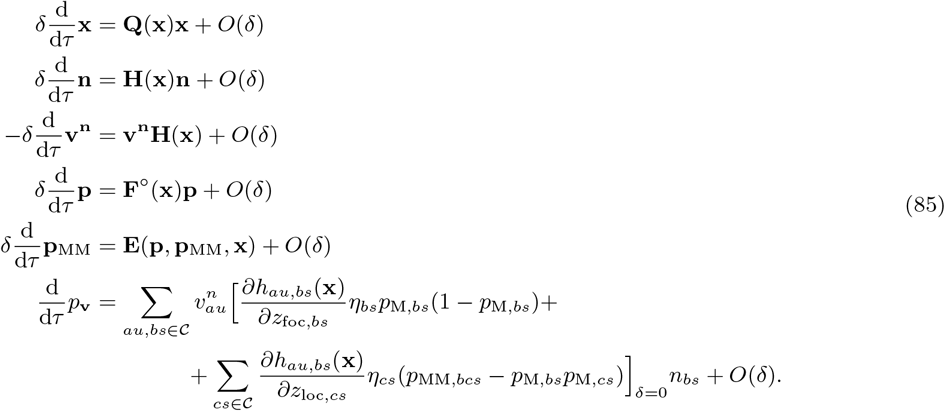

The two systems (84) and (85) are equivalent, the only difference is the notation. Next we study the fast and slow subsystems of (84) and (85) by setting *δ* = 0, respectively, and then in Section 7 we will join the two subsystems together.

### 6.1 Fast subsystem and the critical manifold

We obtain the fast subsystem by setting *δ* = 0 in (84), which yields

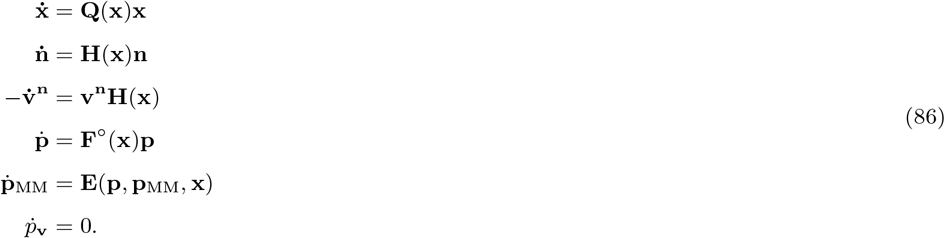

The fast variables (**x, n, v**^**n**^, **p, p**_MM_) thus change in the fast subsystem while the purely slow variable *p*_**v**_ stays constant. More specifically, whenever 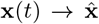 as *t* → ∞ (Section 3.5), then also 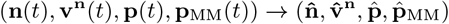). Recall that the vectors 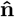 and 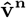 describe the unique steady state of the size and class structure (Section 3.5), and 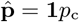 and 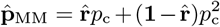 give its genetic structure at the steady state of the fast subsystem. Because *p*_**v**_(*t*) = *p*_c_ for all *t* (Section 5.1), the variable *p*_**v**_ is a constant in the fast subsystem (86) and we can identify *p*_c_ with *p*_**v**_. Thus, as *t* → ∞, the general solution of (**x, n, v**^**n**^, **p, p**_MM_, *p*_**v**_) converges to a curve 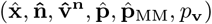 that is parametrized by *p*_**v**_ ∈ [0, 1], and a specific ‘initial value problem’ with an initial condition (**x**(*t*_0_), **p**(*t*_0_)) for some initial time *t*_0_ converges to a single point on this curve.

We collect all the steady states of the fast subsystem into a single set

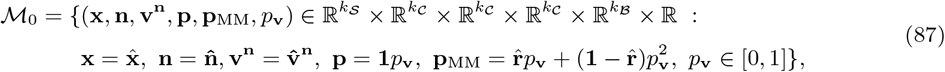

which defines the critical manifold of the system (e.g., Kuehn, 2015 for general considerations and Priklopil and Lehmann, 2020 in the context of the “invasion implies substitution”-principle). The critical manifold thus gives the steady states of the *general* solution of (86) and is a curve consisting of an infinite number of steady states (equilibria) parametrized by *p*_**v**_ ∈ [0, 1] (see Figure 2, panels (a)-(c) for a graphical representation in). Because 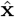 is assumed to be hyperbolically stable (Section 3.5) the critical manifold is said to be a normally hyperbolically stable invariant manifold. For such manifolds the dynamics near the manifold is dominated by the hyperbolicity condition (i.e. in the normal direction to the manifold) while the fast dynamics on the manifold is constant (in the fast subsystem). Indeed, all the variables are constant on ℳ_0_ and because *p*_**v**_ is defined on an interval [0, 1] the manifold is a 1-dimensional manifold in a (3*k*_𝒞_ + *k*_𝒮_ + *k*_ℬ_ + 1)-dimensional space and can be seen as the state space for the pure slow dynamics where *δ* = 0 in (85) (studied in detail in Section 6.2). This manifold plays an important role in the proof of the “invasion implies substitution”-principle: in the fast-slow mutant-resident system where *δ* is small but non-zero the critical manifold persists as a so-called slow manifold (Fenichel, 1979), and, the dynamics of the fast-slow mutant-resident system (84)-(85) on and near the slow-manifold is equivalent to the dynamics of *p*_**v**_ on and near ℳ_0_ in the slow subsystem (Section 6.2).

**Figure 2:**
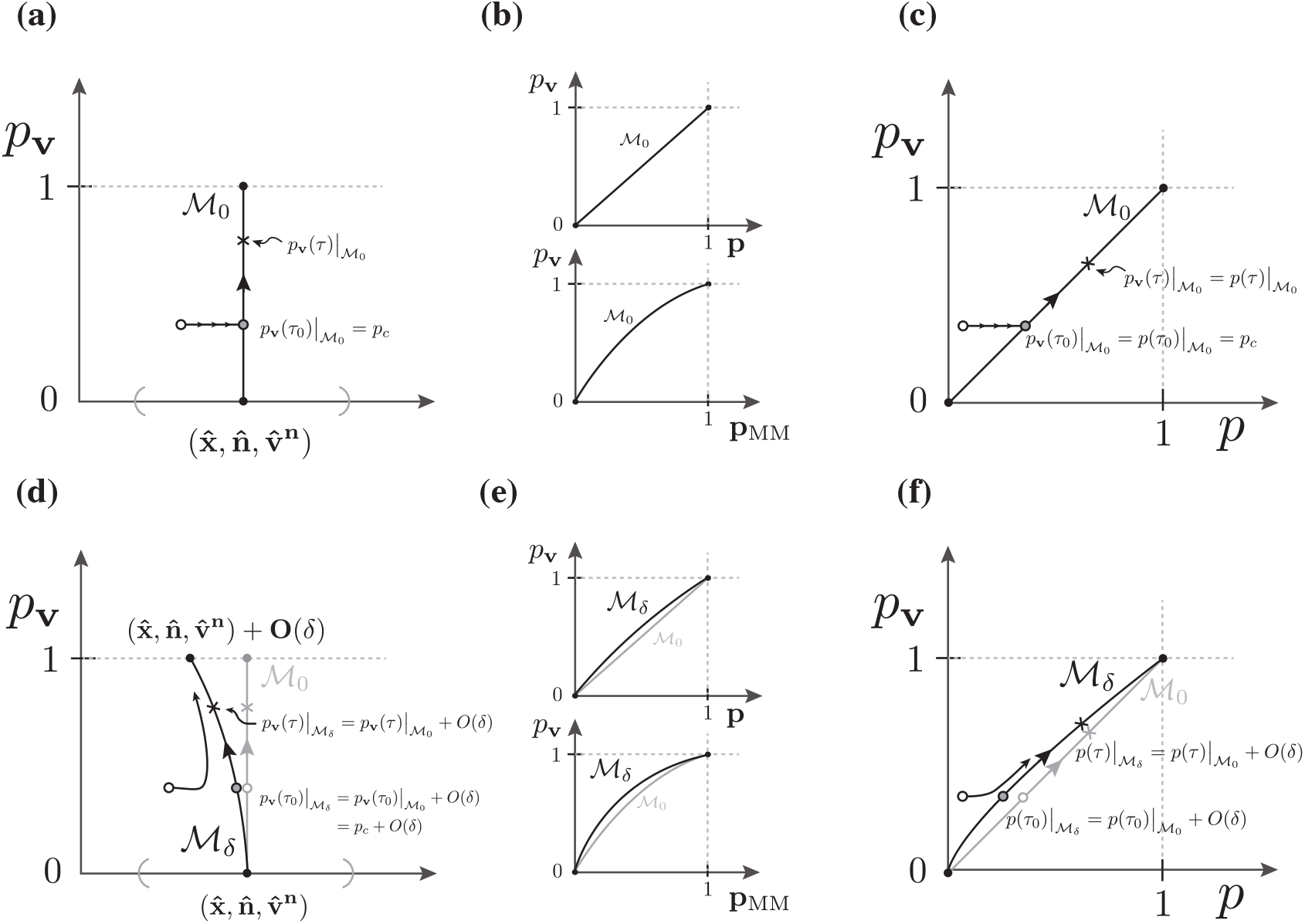
Panels (a)-(c) depict the fast and slow subsystems and the critical manifold ℳ_0_ (86)-(87) (where *δ* = 0, Sections 6.1-6.2), and, panels (d)-(f) depict their singular perturbations and the slow manifold ℳ_*δ*_ (where *δ* small but non-zero, Section 7). **(a)** The phase-plane for the fast variables (**x, n, v**^**n**^) (the “*x*-axis”) and the slow variable *p*_**v**_. In the fast subsystem (Section 6.1) the slow variable *p*_**v**_ stays constant while the fast variables converge to 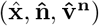 whenever **x**(*t*_0_) is in the neighborhood (indicated by gray semi-circles) of 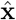 for some initial time *t*_0_. A specific solution for the fast subsystem is depicted with small arrows and where the empty circle shows the initial condition for *t*_0_. The slow subsystem (Section 6.2) is constrained on ℳ_0_, and a particular solution is constructed by taking the steady state of the fast subsystem (where *t* → ∞) as the initial condition for the slow subsystem 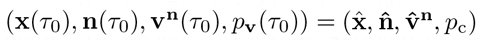. In this example *p*_**v**_(*τ*) → 1 as *τ* → ∞. **(b)** The shape of ℳ_0_ on the (**p**, *p*_**v**_)-plane where 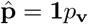 (top panel) and (**p**_MM_, *p*_**v**_)-plane where 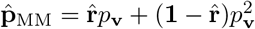 (bottom panel). Note that the shape of ℳ_0_ in the bottom panel depends on the steady state for relatedness 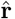. **(c)** Phase-plane for the fast (and slow) arithmetic mutant frequency *p* (55) and the slow reproductive value weighted mutant frequency *p*_**v**_ (the notation is identical to panel (a)). Note that *p* = *p*_**v**_ on ℳ_0_ and that their dynamics on ℳ_0_ is as given in (88) and (91). **(d)** The phase plane for (**x, n, v**^**n**^, *p*_**v**_) as a singular perturbation of (a) where, in particular, the slow manifold ℳ_*δ*_ is *O*(*δ*)-close to ℳ_0_ as is the initial condition (gray circle) and the solution on ℳ_*δ*_. Moreover, because ℳ_*δ*_ is a stable manifold, the solutions in the neighborhood of ℳ_*δ*_ approach the solutions on M_*δ*_. **(e)** The shape of ℳ_*δ*_ as a perturbation of ℳ_0_ discussed in (b). **(f)** The phase plane for (*p, p*_**v**_) as a singular perturbation of (c). Note that the solutions *p*(*τ*)| ℳ_*δ*_ and *p*_**v**_(*τ*) ℳ_*δ*_ are *O*(*δ*)-distance away from each other because ℳ_*δ*_ lies off-diagonal. The interpretation of the solutions are identical to (d).

### 6.2 Slow subsystem on the critical manifold

We obtain the slow subsystem by setting *δ* = 0 in (85). However, motivated by Section 6.1, we are in fact not interested in every possible slow subsystem of (85), only the one associated to 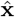 and the critical manifold ℳ_0_ (87). Indeed, there may be other steady states satisfying **0** = **Q**(**x**)**x** and hence other critical manifolds and slow subsystems. The slow subsystem of interest is then given jointly by the critical manifold ℳ_0_ in (87) and the dynamics of the weighted mutant frequency *p*_**v**_ restricted to this manifold,

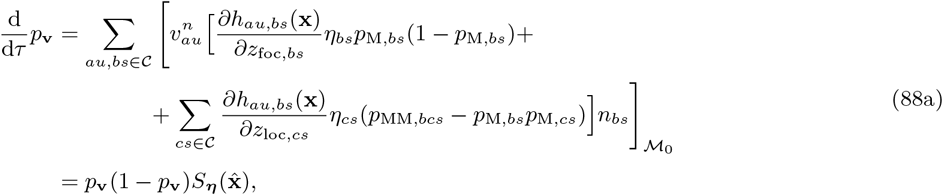

where

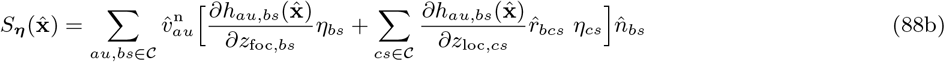

is evaluated at the critical manifold ℳ_0_. To obtain equation (88) we have taken the limits *δ* → 0 and *t* → ∞ and recall that throughout we have assumed that the total population size is infinity. Equation (88) can thus be seen as the first moment of change in the reproductive value weighted allele frequency of the diffusion approximation for slow subsystems (Ethier and Nagylaki, 1980, 1988).

### 6.3 Solution of the slow subsystem

Given that the dynamics of the weighted mutant frequency *p*_**v**_ can be solved from a single ODE given in (88), we can express this in the context of an initial value problem as follows. First, recall that the fast subsystem (86) converges to ℳ_0_ as *t* → ∞, and that the exact point on ℳ_0_ that the fast subsystem converges to depends on the given initial condition (**x**(*t*_0_), **p**(*t*_0_)) which determines *p*_**v**_ = *p*_c_ ∈ [0, 1] that parametrizes ℳ_0_ (Section 6.1). We will assume that this point is an initial condition for the slow subsystem, and to this end we define *p*_**v**_(*τ*_0_) = *p*_c_ for some initial slow time *τ*_0_, and let

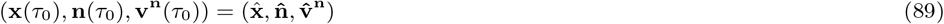

and

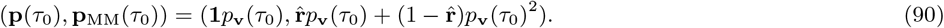

Note that the dependency of (89)-(90) on (**x**(*t*_0_), **p**(*t*_0_))) comes only through (90). Then, in slow time *τ*, the weighted mutant frequency *p*_**v**_(*τ*) changes according to (88) for all *τ* ≥ *τ*_0_, and for the initial condition (89)-(90) its solution is

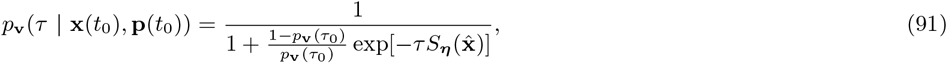

where 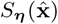 is given in (88b) and where we have indicated that the initial condition *p*_**v**_(*τ*_0_) = *p*_c_ depends on (**x**(*t*_0_), **p**(*t*_0_)).

It is important to note, that while 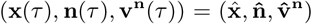 stay constant for all *τ* ≥ *τ*_0_, the genetic structure changes according to **p**(*τ*) = **1***p*_**v**_(*τ*) and 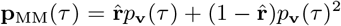 for all *τ* ≥ *τ*_0_ and where *p*_**v**_(*τ*) = *p*_**v**_(*τ* | **x**(*t*_0_), **p**(*t*_0_)) is the solution in (91). Because **p** and **p**_MM_ change also in the fast subsystem (Section 6.1), they are both fast *and* slow variables. Moreover, because every average mutant frequency *p*_***ϕ***_ in (65) in the fast subsystem converges to *p*_c_ as *t* → ∞ (Section 5.1), we will also define

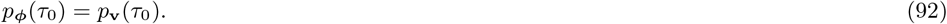

Then, *p*_***ϕ***_(*τ*) = *p*_**v**_(*τ*) for all *τ* ≥ *τ*_0_, and the solution of *p*_***ϕ***_ and thus also the solution of the arithmetic mean mutant frequency *p* in (55), is identical to the solution (91). See Figure 2, panels (a)-(c) for a graphical representation.

## 7 Invasion implies substitution

We are now ready to state and prove the “invasion implies substitution”-principle for the metacommunity model.

### Invasion implies substitution – principle

*Suppose that resident individuals express phenotype* ***z*** ∈ Ƶ *and that the resident dynamics of the metacommunity is given as in* (10). *Suppose further that the metacommunity is near a hyperbolically stable steady state* 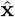 *that satisfies* 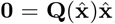, *and that a mutation generates a mutant phenotype* ***z*** + *δ****η*** *where* ***η*** *gives the direction of the deviation and δ characterizes the distance between mutant and resident phenotypes. Then, for sufficiently small δ and/or large time t* ≥ *t*_0_ *where t*_0_ *is some initial time, the dynamics of the arithmetic mean mutant frequency p can be approximated as*

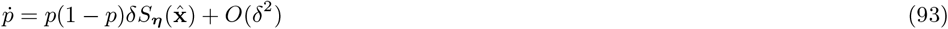

*with solution*

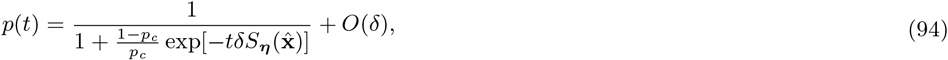

*where*

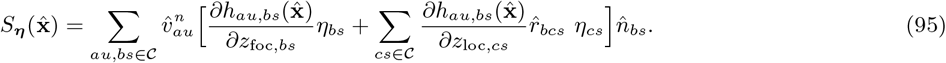

*The constant p*_*c*_ *in* (94) *is the asymptotic mean mutant frequency of the fast subsystem where δ* = 0 *and it depends on the initial condition* (**x**(*t*_0_), **p**(*t*_0_)) *where* **p**(*t*_0_) *is the initial distribution of within-class mutant frequencies. In* 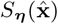, *the density distribution vector* 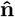 *and the reproductive value vector* 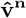 *are the right and left dominant eigenvectors of* 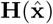 *with a scaling* 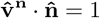 *where* 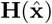 *is the resident fitness matrix in* (15). *The relatedness vector* 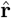 *satisfies* 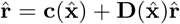 *where* 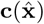 *is a vector giving the probabilities that a pair of individuals have a common parent and* 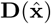 *is a matrix giving the probabilities that this pair has two distinct parents within the same group* *(Section 3.4.1)*.

### 7.1 Proof of the principle

Here we give a poof of the “invasion implies substitution”-principle stated above. First, recall that in Section 4 we set up a mutant-resident metacommunity model for arbitrary *δ*, and in Section 5 we showed that in order to study the spread of a mutant allele for small *δ* we can study the fast-slow dynamics of five fast variables (**x, n, v**^**n**^, **p, p**_MM_) dominated by *O*(1) and a single slow variable *p*_**v**_ dominated by *O*(*δ*). To do this, in Section 6 we set up a fast-slow mutant-resident system for (**x, n, v**^**n**^, **p, p**_MM_, *p*_**v**_) and then analyzed its fast and slow subsystems where *δ* = 0. In the fast subsystem (Section 6.1), we found that whenever **x** converges to 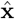, all the remaining fast variables (**n, v**^**n**^, **p, p**_MM_) of interest converge to the critical manifold ℳ_0_ at which 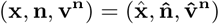 are isolated steady state points and 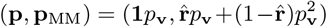 take values on a curve parametrized by *p*_**v**_ ∈ [0, 1] which is a constant in the fast subsystem (and hence is on ℳ_0_ by definition). The exact point *p*_**v**_ = *p*_c_ ∈ [0, 1] on ℳ_0_ that the fast subsystem converges to depends on the initial condition (**x**(*t*_0_), **p**(*t*_0_)) where **x**(*t*_0_) is chosen sufficiently close to 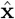.

In the slow subsystem (Section 6.2), we found that the slow variable *p*_**v**_ changes according to an ODE given in (88) and when coupled with an initial condition its solution is given by (91). Consequently, the fast variables 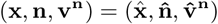 stay constant in the slow subsystem and are thus purely fast variables, and the two moments of allelic states 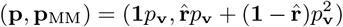 change due to *p*_**v**_ and are hence both fast and slow variables. Next, we perturb the fast and slow subsystems using classical results from geometric singular perturbation theory (Fenichel, 1979; Hek, 2010; Kuehn, 2015; Priklopil and Lehmann, 2020), which proves the “invasion implies substitution”-principle for small but nonzero *δ*. To this end, we proceed in two steps. In **step 1** we apply ‘Fenichel’s first theorem’ (e.g. Priklopil and Lehmann, 2020, Appendix A.4., Fenichel’s invariant manifold theorem 1) saying that because ℳ_0_ is a normally hyperbolic invariant manifold, for small but non-zero *δ* there exists a so-called slow manifold ℳ_*δ*_ that is **(i)** *O*(*δ*)-close and diffeomorphic to ℳ_0_, and, **(ii)** invariant under the fast-slow mutant resident dynamics (84)-(85). Property **(i)** says that for any point on ℳ_0_ there is a point on ℳ_*δ*_ that is *O*(*δ*)-close. We will state this in terms of an initial condition of the slow subsystem as

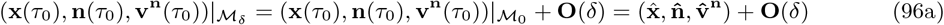

and

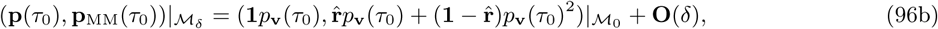

and which is valid for any initial condition (i.e., any point on ℳ_0_) and where **O**(*δ*) is a vector of *O*(*δ*). The property **(ii)** says that for all *τ* ≥ *τ*_0_, the solutions starting on ℳ_*δ*_ will remain on ℳ_*δ*_. Importantly, the two properties **(i)** and **(ii)** together imply that for all *τ* ≥ *τ*_0_ the ODE’s (or vector fields) and their solutions on ℳ_*δ*_ are small perturbation of the ODE’s (vector fields) and solutions on ℳ_0_. More precisely, because the dynamics of *p*_**v**_(*τ*) on ℳ_0_ denoted as *p*_**v**_(*τ*)|_M0_ can be written as an ODE (88) and when coupled with an initial condition its solution is given as in (91) for all *τ* ≥ *τ*_0_ (Section 6.2), the dynamics of *p*_**v**_ on ℳ_*δ*_ can be written as an ODE

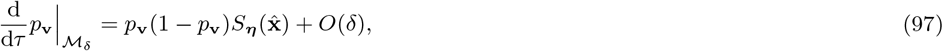

and when coupled with an initial condition given in (96), its solution is

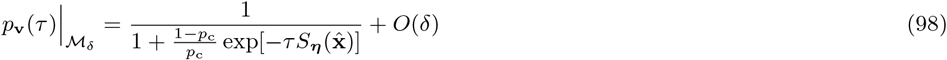

where *p*_c_ = *p*_**v**_(*τ*_0_) is the initial weighted mutant frequency in the slow subsystem. The equations (97)-(98) are the first order approximations of the dynamics on ℳ_*δ*_.

So far we focused on the dynamcis of *p*_**v**_(*τ*). But since on ℳ_0_ we have *p*_**v**_(*τ*) = *p*_***ϕ***_(*τ*) for all *τ* ≥ *τ*_0_ (Section 6.3), and because *δ* is nonzero in (97)-(98), we can re-write (97)-(98) in fast time *t* = *τ/δ* and in terms of *p*_***ϕ***_ as

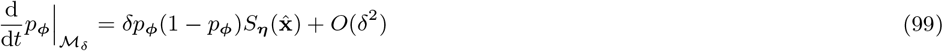

and

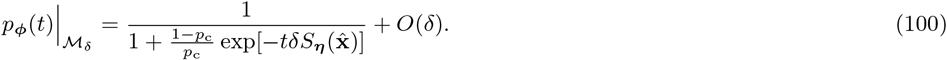

Equations (97)-(100) thus establish the first order approximation for any average mutant frequency in the fast-slow mutant resident system (84)-(85) restricted to the slow manifold ℳ_*δ*_. We note that while the solutions *p*_**v**_ and *p*_***ϕ***_ on ℳ_*δ*_ are both *O*(*δ*)-distance away from their solutions on ℳ_0_, they are also *O*(*δ*)-distance away from each other (Figure 2 panel (f), see further discussion below).

It now remains to investigate whether the first order approximations (97)-(100) on ℳ_*δ*_ are generic in the sense that any other solution nearby is a small perturbation away, which is a necessary requirement for us to be able to choose initial conditions away from ℳ_*δ*_. This leads to **step 2** of the proof in which we apply Fenichel’s results on the persistence of the stability properties of manifolds (e.g. Priklopil and Lehmann, 2020, Appendix A.4., Fenichel’s invariant manifold theorems 2 and 3). These results say that because ℳ_0_ is attracting so is ℳ_*δ*_, and moreover, all the variables near ℳ_*δ*_ converge to ℳ_*δ*_ exponentially fast. The exact rates can be estimated as given in Priklopil and Lehmann (2020, Appendix A.4., Corollaries 1 and 2). Therefore, a solution *p*_***ϕ***_(*t*) with an initial condition *p*_***ϕ***_(*t*_0_) near ℳ_*δ*_, for some initial time *t*_0_, will become arbitrarily close to (99)-(100) in finite time that depends on *δ*. See Figure 2 panels (d)-(f) for a graphical representation of the proof.□

Using the weighted mutant frequency *p*_**v**_ and geometric singular perturbation theory (Fenichel, 1979; Hek, 2010; Kuehn, 2015), we showed that the “invasion implies substitution”-principle holds for our metacommunity model, saying that the dynamics of any average population-wide mutant frequency *p*_***ϕ***_ can in fact be approximated for sufficiently small *δ* and/or large fast time *t* by a *single* scalar valued dynamical equation (93)-(95). Moreover, the rate at which change in mutant frequency occurs in the ‘normal’ population dynamical time *t* is characterized by 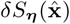 giving the number of additional mutant alleles produced per unit of time by the focal mutant allele (see further discussion in Section 7.2). This can be expressed purely in terms of quantities obtained from the resident metacommunity dynamics evaluated at their steady states, and are thus constant over the relevant timescales. We note that because the rate of mutant frequency change is measured per unit of time, this rate is of order *O*(*δ*) slower in fast time *t* than in slow time *τ* because 1*/t* = *δ/τ* (compare e.g. (97) and (99)), and thus 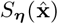 characterized the rate of mutant frequency change in the slow evolutionary time *τ*. In contrast, the solution of the average mutant frequency *p*_***ϕ***_ of the fast-slow system where *δ* is small is *O*(*δ*)-distance away from the solution of the slow subsystem (where *δ* = 0), and this distance is independent of the timescale (compare e.g. (98) and (100)). This implies that the per unit time change in the arithmetic mean mutant frequency is *δ*-distance away from the asymptotic change in the reproductive value weighted frequency. Moreover, as *δ* approaches 0 but remains non-zero, the slow subsystem (91) gives an increasingly good approximation to (94).

### 7.2 Inclusive fitness effect and singular points

An alternative representation of the directional selection coefficient 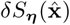 as characterised in (95) can be obtained by considering a class-frequency vector **u** = (*u*_*as*_)_*as*∈𝒞_ = **n***/n* where *n* = ∑_*as*∈𝒞_ *n*_*as*_ is the total density of the focal species. Because each individual can be identified with the allele it carries, we will interpret *n* as the total number of mutant and resident alleles in the focal species and **u** as the distribution of the *context* they find themselves in (Kirkpatrick et al., 2002), and recall that the distribution of mutants and residents is given by **p**. In other words, given we randomly sample a carrier of the mutant allele (or resident allele) at the steady state for a process where *δ* = 0, _û*as*_ is the probability that this allele finds itself in class *as* ∈ 𝒞. Then, by re-scaling the critical manifold (87) as 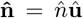 and 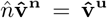 such that 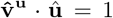 where **v**^**u**^ is the adjoint variable associated to **u** (and corresponds to the standard scaling of reproductive value, e.g., Taylor, 1990), the directional selection coefficient 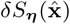 characterised in (95) can be written and interpreted as given in (2) (by adding a superscript **u** to **v**). Note that in (2) we write *S*_***η***_ (***z***) thus using ***z*** instead of 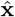 as the main argument because it is commonly used as such to study phenotype evolution and we shall follow this notational change henceforth. We emphasise that under both scalings, (2) and (95), the directional selection coefficient measures the ‘additional’ mutant alleles produced by a randomly sampled carrier of the mutant allele, i.e., a focal mutant individual. This is true in particular in the second summation in (2) and (95) where the contribution of group members is accounted for.

We can now further change the perspective on the contribution of group members by swapping indices in the second summation in (2), and using *u*_*cs*_ = (*N*_*cs*_*/N*_*bs*_)*u*_*bs*_ for all *bs, cs* ∈ 𝒞, we then obtain the representation

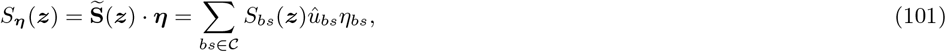

with 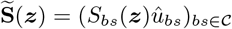 where

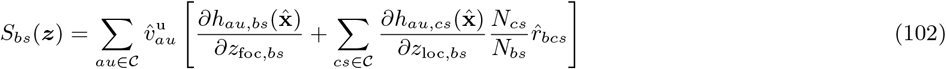

is the selection pressure on phenotypic component *z*_*bs*_ ∈ *Ƶ*_𝒞_. Similarly to (2) and (95), the first term in the brackets 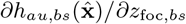, when multiplied by *δ*, gives the ‘additional’ number of gene copies (or offspring number owing to asexual reproduction) in class *au* ∈ 𝒞 produced per unit of time by an individual of class *bs* ∈ 𝒞 as a consequence of it bearing the mutant instead of the resident allele. In contrast to (2) and (95), however, the second term 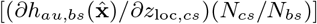 in the second summation, when multiplied by *δ*, gives the ‘additional’ number of class *au* ∈ 𝒞 gene copies produced per unit of time by all *non-focal* local individuals of class *cs* ∈ 𝒞 (group members of the focal individual) as a consequence of a focal individual of class *bs* ∈ 𝒞 bearing the mutant instead of the resident allele. The weight 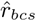 ensures that we only count those non-focal individuals *cs* ∈ 𝒞 who have a common ancestor with the focal individual *bs* ∈ 𝒞 and are thus likewise mutants. Each such additional mutant gene copy in class *au* ∈ 𝒞 is then weighted with its reproductive value 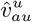 that gives the expected asymptotic *number* of descendants, and we finally sum over all classes of the offspring *au* ∈ 𝒞. The directional selection coefficient *δS*_***η***_ (***z***) thus gives the number of additional mutant alleles produced either directly (the first term in (102)) or indirectly (via related individuals, the second term in (102)) per unit of time by a focal mutant allele that is distributed according to _û_. That is, in (101)-(102), the individual fitness ‘belongs’ to the focal individual in the first term in (102) (as in (2) and (95)) whereas in the second term in (102) it belongs to non-focal but related individuals (in contrast to (2) and (95)) that are affected by the focal’s trait(s), whereby the focal individual can be taken as the actor of all fitness changes as according to inclusive fitness theory (Hamilton, 1970). Hence, (101) separates mutational step size from structure of selection by associating all fitness consequences of an allele that would induce an unilateral phenotypic change *η*_*bs*_ into *S*_*bs*_(***z***) for all *bs* ∈ 𝒞 in (102).

It then follows from (101) that a singular point ***z***^*^ ∈ *Ƶ* needs to satisfy

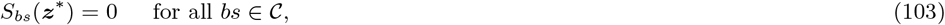

since at a singular point no unilateral deviation in phenotype should result in a change in fitness, that is, (101) needs to be 0 for all ***η***. Whether such a singular point will be approached by gradual evolution from within its neighbourhood hinges on the concept of convergence stability (Eshel, 1983; Lessard, 1990; Christiansen, 1991; Leimar, 2009). However, because convergence stability along with other properties of singular points (e.g. evolutionary branching Geritz et al., 1998; Ajar, 2003) is a ‘second order’ property we leave such further analysis for future work. Here we simply state that while in the context of multidimensional phenotypic evolution the direction of change of phenotype can depend on the mutation process, the covariance of the class-specific mutational increments are unlikely to interfere with the convergence stability of singular points when the mutation matrix is positive definite (Leimar, 2005, 2009).

## 8 Discussion

We have proved the “invasion implies substitution”-principle ((93)-(95), Section 7) for the evolution of a multidimensional quantitative trait expressed by the individuals of a focal species that is physiologically class structured e.g. by age, size and stage, and that resides in a metacommunity with finite local community size where reproduction, dispersal, survival, and development are stochastic and frequency- and density-dependent.

### 8.1 Evolutionary dynamics across multiple timescales

The evolutionary dynamics discussed in this paper are driven by three qualitatively different evolutionary forces (processes): mutation, selection and class-transmission. Mutation-driven changes in the metacommunity are caused by the apparently random modification of the expression of the phenotype. And while selection is classically driven by the differential survival and reproduction due to differences in phenotypes among individuals, class-transmission is driven by the differential survival and reproduction due to individuals residing in different classes. Mutant frequency can thus change not only due to selection but *also* transmission, even in the absence of selection (see (52)). In the following we will examine all these processes in detail and how they contribute to the adaptive dynamics of a quantitative trait.

In the mutant-resident metacommunity model with arbitrary phenotypic effects (Section 4), both selection and class-transmission are arbitrarily strong and intertwined. They together drive the change in mutant allele frequency at all times, a change that can involve density-dependent and frequency-dependent dynamics potentially leading to allelic coexistence (polymorphism). In Section 5, however, we showed (as expected) that under the assumption (i) of phenotypic closeness, the effect of selection on the mutant frequency dynamics is much weaker than the effect of class-transmission. Hence, the large system of mutant frequency dynamics can be partitioned into two simpler limiting subsystems (Section 6), one that is driven only by class-transmission on a fast population dynamical timescale (fast subsystem, Section 6.1) and the other driven only by selection on a slower micro-evolutionary timescale (slow subsystem, Section 6.2). One can further analyze these subsystems in isolation and join them together using geometric singular perturbation methods (Fenichel, 1979; Hek, 2010; Kuehn, 2015) eventually leading to the “invasion implies substitution”-principle (93)-(95) (Section 7). In order to follow these steps, however, we had to overcome the difficulty that in class structured populations there does not exist *a priori* a purely slow variable driven by selection only and that can be used as a proxy for mutant frequency (e.g. Rousset, 2004; Priklopil and Lehmann, 2020). Such a variable can nevertheless be constructed by weighting the distribution of mutants by reproductive values (Taylor, 1990; Rousset, 2004; Grafen, 2015; Lion, 2018b; Priklopil and Lehmann, 2020), because by definition such weights scale away any changes due to class-transmission. Hence, the assumption of small phenotypic differences and the use of the weighted frequency *together* lead to the separation of timescales and the consequent separation of evolutionary dynamics into frequency changes caused by class-transmission and selection, which in turn underlies arithmetic mean allele frequency change (93)-(95) whose sign is independent of frequency. Our result thus extends earlier work whose representation of (1) was given in terms of a reproductive value weighted mutant frequency and not the arithmetic mean (Roze and Rousset, 2004; Rousset and Ronce, 2004; Rousset, 2004; Van Cleve et al., 2010; Priklopil and Lehmann, 2020).

We must stress that the arithmetic mean allele frequency change (93)-(95) holds not only in the limiting slow subsystem where *δ* → 0 and where the population dynamical time *t* → ∞ (Section 6.2), but also for small but non-zero *δ* in the ‘normal’ population dynamical time *t*^*^ ≤ *t* ≤ ∞ where *t*^*^ is some finite time after which (93)-(95) hold (Section 7). In this respect, our results complement earlier work on the mutant allele dynamics in slow subsystems (e.g. Wakeley, 2003; Roze and Rousset, 2003; Rousset, 2004; Wakeley and Takahashi, 2004; Lessard, 2009, and see for well-mixed but age and/or class structured populations Lessard and Soares, 2018; Soares and Lessard, 2019, 2020). In this work, the strength of selection *δ* is inversely proportional to the system size that tends to infinity while *δ* → 0 but keeping their product finite, and it invokes a two timescale method of Ethier and Nagylaki (1980). Our result (1)-(2) for small but nonzero *δ* holds whenever one can show that the metacommunity is ‘structurally stable’ in the sense that the steady state of the metacommunity and its stability properties persist under the small perturbation caused by the invasion of the mutant, that is, when *δ* deviates from 0. In the present metacommunity model, this is guaranteed by the assumption of hyperbolicity (ii), and jointly with the assumption on phenotypic similarity (i) and the weighted frequency *p*_**v**_ fully justify the “invasion implies substitution”-principle (93)-(95). Recurrent invasion-substitution events thus cause the seemingly gradual changes in the structure of the metacommunity and the gradual evolution of the phenotype of interest.

The final question is whether and how is the process of gradual evolution affected by the mutation processes. For scalar-valued phenotypes, recurrent invasion-substitution events will always either increase or decrease the phenotypic value until the phenotype becomes a singular point and the same should hold for multidimensional traits under mild assumptions on the mutation matrix when the singular point is convergence stable (see section 7.2). Interestingly, a mutation rate that results in several competing mutations in the metacommunity does not interfere with the qualitative nature of this gradual process. This is because any finite number of overlapping mutations perturb the structure of the resident species only to order *O*(*δ*) (Meszéna et al., 2005). Therefore, if the mutation of interest results in a positive selection coefficient *δS*_***η***_ (**z**) in a resident metacommunity, the selection coefficient remains positive upon the arrival of any new mutations, provided the assumption (i) for small phenotypic effects holds. Naturally, this is also true for the other ‘overlapping’ mutations: *any* mutation that results in a positive selection coefficient *δS*_***η***_ (**z**) will go to fixation upon a successful invasion event. A mutation rate that generates several overlapping mutations thus only affects the rate of evolution but not its direction. Furthermore, it is important to emphasise that the “invasion implies substitution”-principle (93)-(95) is valid also for polymorphic resident metacommunities (Priklopil and Lehmann, 2020). In such a case the “invasion implies substitution”-principle holds along each “morph” of the polymorphic resident species requiring as many directional selection coefficients as there are morphs. Therefore, provided we stay away from singular points and mutations have small phenotypic effects, the “invasion implies substitution”-principle (1)-(2) gives a complete description of the gradual evolutionary process.

### 8.2 Evolutionary dynamics across multiple fluctuations

Because we have formulated the metacommunity dynamics as a stochastic processes that can depend on the state of the group and the metacommunity at large, our model captures frequency- and density-dependent environmental feedbacks both at the group-as well as metacommunity-level. Our model also applies to situations where reproduction, dispersal, survival and physiological development depends on locally fluctuating external factors, defined as any factors that are not contained in the environmental feedback, and can thus generate the effect of environmental stochasticity on a local (group) scale and whenever the external factors can be expressed as discrete variables. Extending our model to capture globally fluctuating external factors would require to introduce an additional “environmental” state space *ε* whose elements would affect all quantities used in this paper but not change their functional form, that is, all rates used in this paper would also be functions of ε (in the context of this present paper but for well-mixed populations see a recent approach in Cai and Geritz, 2020). Yet, we expect that by taking an appropriate average over the ergodic path of environmental states including such environmental change is unlikely to change the generic shape of the selection coefficient, a point illustrated by Lenormand et al. (2009), proven more generally by Lion (2018b, eq. 23) for panmixia, and implied by the analysis of Svardal et al. (2015, eq. B.8) for spatial structure. Moreover, in metacommunity models where the total size of the focal species is finite, our dynamical equations can in principle be used to construct a diffusion approximation with two timescales to analyze the effect of genetic drift on the evolutionary dynamics (e.g., Wakeley, 2003; Roze and Rousset, 2004; Rousset, 2004; Soares and Lessard, 2020). Here, the infinitesimal mean of the diffusion is given by the mean change (88) and the infinitesimal variance is obtainable from the recurrence equations for relatedness (Rousset, 2004, chapter 9-11). Finally, we formulated our model only in the context of haploid reproduction. Still, allowing for diploidy and sexual reproduction under additive gene action within individuals should only require changing the scaling of the mean change in allele frequency (93) and to interpret relatedness in the directional selection coefficient (95) in a diploid context (as per simpler models with diploidy and class structure, Roze and Rousset, 2004; Van Cleve et al., 2010). In conclusion, while we have left several biological questions to be settled in future work, we believe that the present model contributes to the understanding of adaptive evolution of structured natural populations.

## Acknowledgment

TP was supported by the Swiss NSF grant PP00P3-123344 to LL.

## A Appendix: Individual fitness

### A.1 Resident individual fitness

In this section we detail the steps to obtain individual fitness (16) from the resident group and metacommunity dynamics (10). Using *n*_*au*_ = *N*_*au*_*x*_*u*_, we have

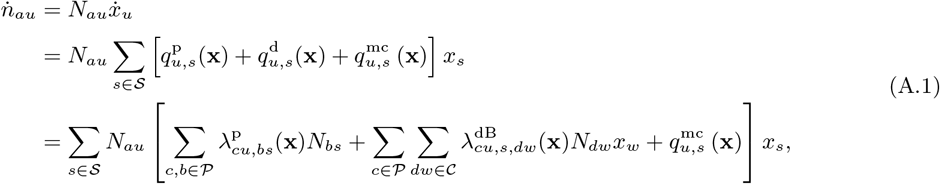

where we have applied Section 3.2. Whenever an individual *a* ∈ 𝒫 is produced or removed from group *s* ∈ 𝒮 so that this group transitions to *u* ∈ 𝒮, the number of *a* ∈ 𝒫 in the group changes and *N*_*au*_ is either equal to *N*_*as*_ + 1 or *N*_*as*_ – 1, otherwise *N*_*au*_ = *N*_*as*_. This motivates us to re-write the above as

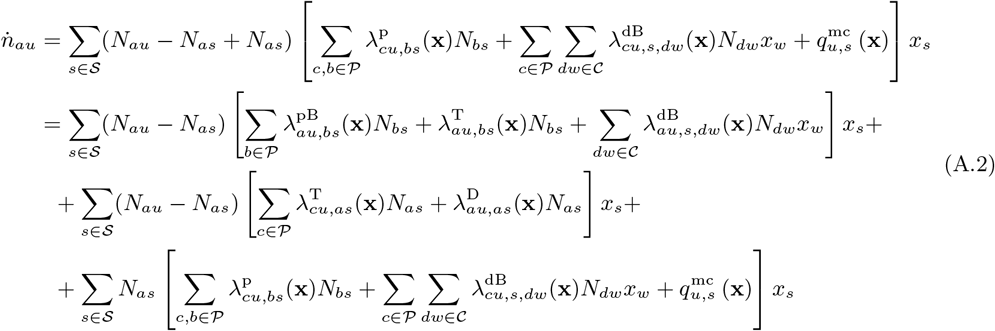

where in the second equality we have applied Sections 3.1-3.2, and by relabeling the indices on the second line in (A.2) we get

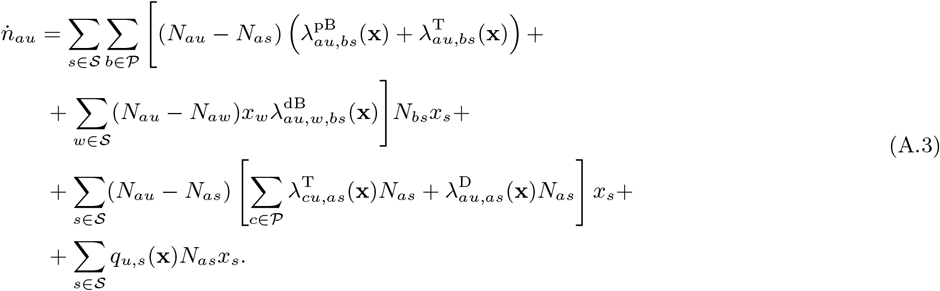

Now, for all *s* ≠ *u* ∈ 𝒮, we have *N*_*au*_ – *N*_*as*_ = 1 and *N*_*au*_ – *N*_*aw*_ = 1 on the first and second line in (A.3), respectively, because an offspring *a* ∈ 𝒫 is produced by birth into a group that transitions to *u* ∈ 𝒮, and *N*_*au*_ – *N*_*as*_ = –1 on the third line in (A.3) because a parent *a* ∈ 𝒫 is removed by physiological transition or death from a group that transitions to *u* ∈ 𝒮. In all other situations where *s* ∈ 𝒮 transitions to *u* ∈ 𝒮 we have *N*_*au*_ = *N*_*as*_. We thus get, after some reorganization,

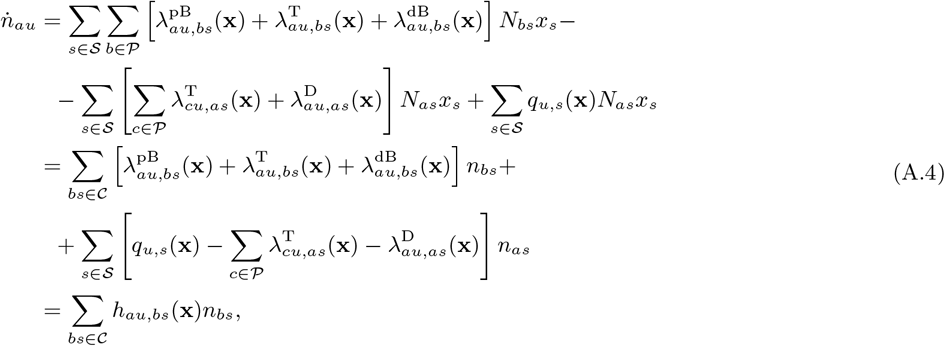

where for *u* ≠ *s* ∈ 𝒮 we can then write

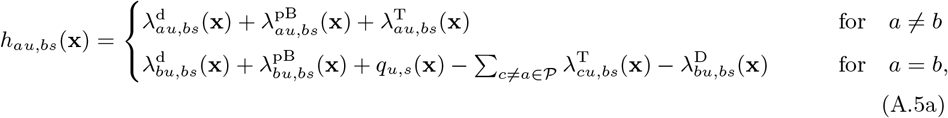

and for *u* = *s* ∈ 𝒮 we have

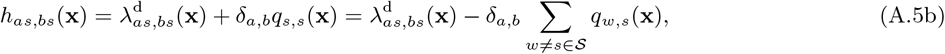

for all *a, b* ∈ 𝒫. Alternatively, we can represent (A.5) with a single equation

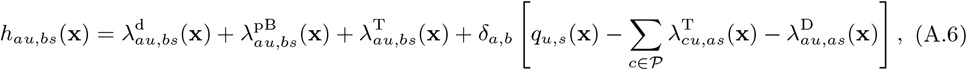

for all *au, bs* ∈ 𝒞, and which is the expression for individual fitness (16) given in Section 3.

#### A.1.1 Individual fitness: connection to previous models and formulations

##### Physiologically unstructured metapopulation model

Here we consider a single physiologically unstructured species and suppose that groups are identical in terms of abiotic factors. The set of g-states can then be represented with 𝒮 = {0, 1, 2, …, *N*_max_} specifying the number of individuals of this focal species inhabiting a group, and the only processes occurring in each group are death, philopatric birth and birth via dispersal, which is conceptually similar to models with local group demographic fluctuations considered previously (Metz and Gyllenberg, 2001; Cadet et al., 2003; Rousset and Ronce, 2004; Lehmann et al., 2006; Alizon and Taylor, 2008; Parvinen, 2013). The individual fitness (16) then reduces to

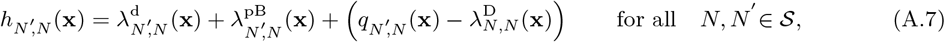

where

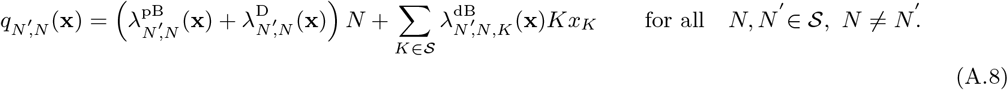

Here,

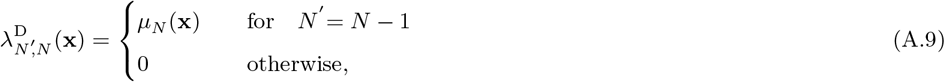

and

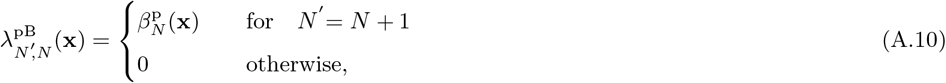

and 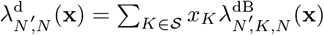 where

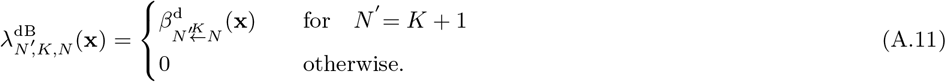

By convention we have *µ*_0_(**x**) = 0, 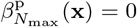 and 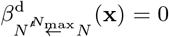. The interpretation of (A.7) is analogous to the one in (16) except that the i-process for physiological transitions as well as all the other processes of non-focal species and abiotic factors are absent. Note that, if we further have 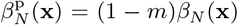 and 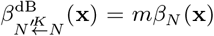 for some migration probability *m* and birth rate *β*_*N*_ (**x**) depending only on the local density of the parent and which is the realm of the aforementioned demographic models, then

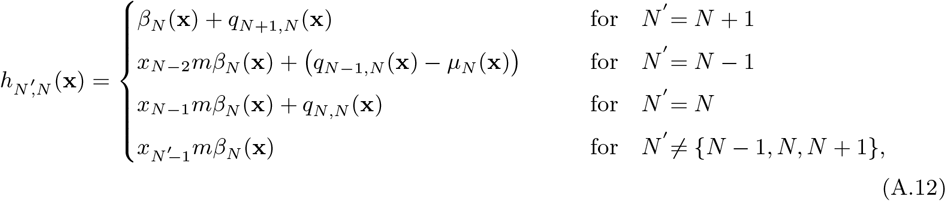

where

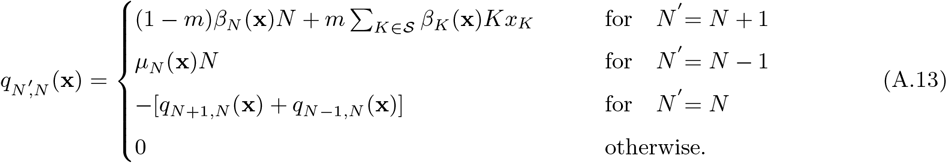

##### Well-mixed physiologically structured population model

Here we reduce our metacommunity model to a well-mixed population model for a single focal species that is physiologically structured. In so doing, we assume that each ‘group’ is inhabited by at most one individual and that individuals reproduce via dispersal into an empty uninhabited group. The set of g-states can be represented with 𝒮 = {0} ⋃ 𝒫 implying that each group is either empty or it is inhabited by a single individual that is in one of the finite number of physiological states. The only processes are birth (via dispersal), physiological transition and death. This model is conceptually similar to the one in Lion (2018a,b) and Priklopil and Lehmann (2020). The individual fitness (16) then reduces to

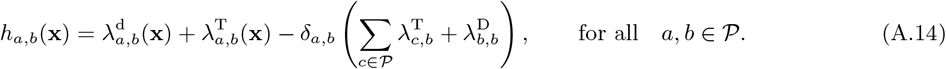

Here, for all *a, b* ∈ 𝒫, we have 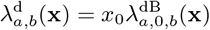 where 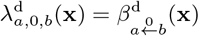, and 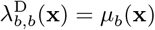 and

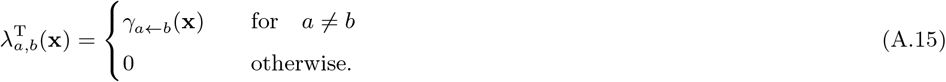

The interpretation of (A.14) is analogous to the one in (16), except that here each ‘group’ can transition from one state to another only due to the i-processes of the focal parent individual *b* ∈ 𝒫. Therefore, the ‘survival’ term in the brackets in (A.14) implies that the focal parent *a* = *b* ∈ 𝒫 has either died and is removed from the group or it physiologically transitions to some other physiological state.

##### Conditional fitness

The concept of individual fitness (16) has been considered in the previous literature for discrete time processes (Lehmann et al., 2016; Ohtsuki et al., 2020) and could generically be called *unconditional* individual fitness because it is evaluated unconditionally on whether or not a group transition occurs. We can distinguish its dispersal and philopatric components, as discussed in Section 3.3, by writing

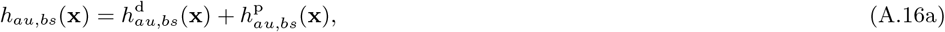

where

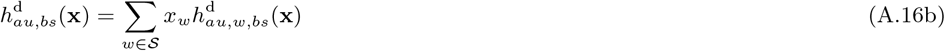

and

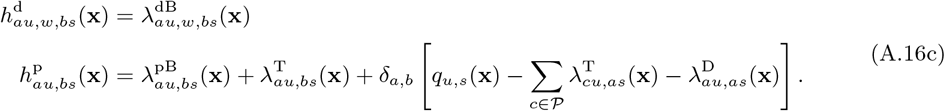

The *conditional* individual fitness (e.g. Rousset, 2004; Rousset and Ronce, 2004,Lehmann and Rousset, 2010 Lehmann et al., 2016, Appendix E) is defined as the fitness of an individual given a group-transition (it is conditional on the realization of the state after the transition), and this can be written as

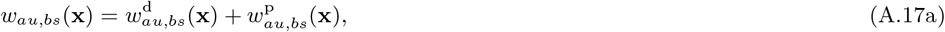

where 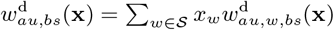 and where

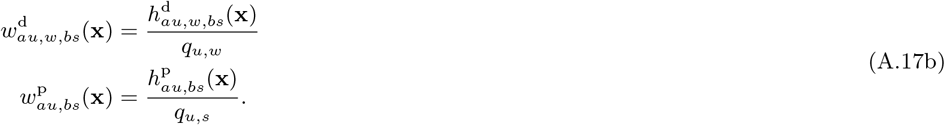

Note that when *a* = *b* ∈ 𝒫, then 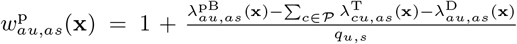 and when *a* ≠ *b* ∈ 𝒫, then 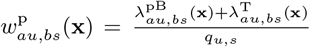. If we now further make the assumptions of the “physiologically unstructured metapopulation model” case where 𝒮 = {0, 1, 2, …, *N*_max_} and use (A.16)–(A.17) along with (A.12)–(A.13) and noting that 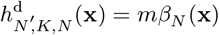 for *N*′ = *K* + 1, then we recover equations (A.43)-(A.48) of Lehmann and Rousset (2010), which hold as the conditional fitnesses for the model of Alizon and Taylor (2008).

### A.2 Mutant-resident individual fitnesses

#### A.2.1 Fitness of mutant and resident individuals

The derivation of fitness functions for mutant-resident dynamics is analogous to the one for resident dynamics (Appendix A, Section A.1). Using 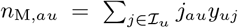 and the fact that upon a transition from *si* ∈ Ω to *uj* ∈ Ω the number of mutants that are added or removed is *j*_*au*_ – *i*_*as*_ for all *a* ∈ 𝒫, we get

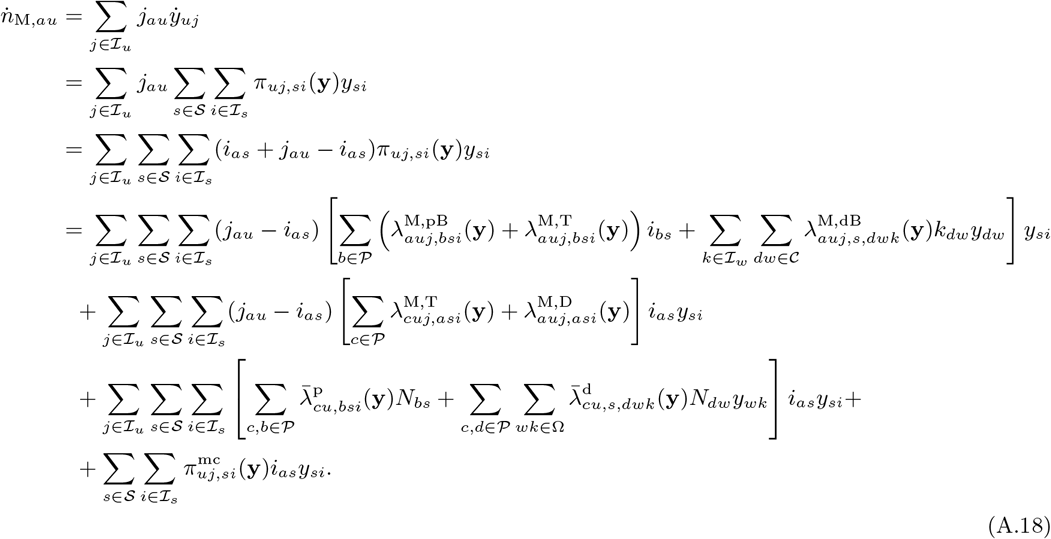

By exchanging the indices and using 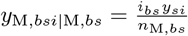, we can re-write the above as

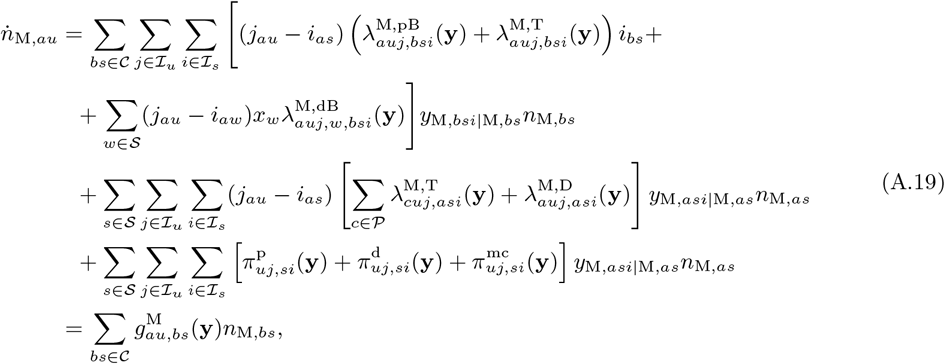

where

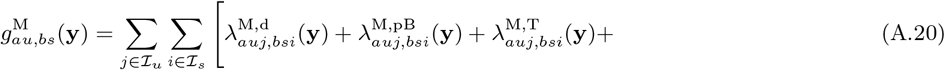

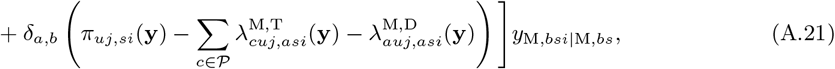

which is the partitioning given in (47).

The derivation of the resident fitness for the mutant-resident dynamics (48) is identical to (A.18)- (A.20), except that one needs to replace the numbers of mutants within a group *i*_*as*_ with resident numbers *N*_*as*_ – *i*_*as*_, for all *as* ∈ 𝒞, class-specific densities for mutants *n*_M,*as*_ with resident densities *n*_R,*as*_, for all *as* ∈ 𝒞, as well as conditional group frequencies *y*_M,*bsi*|M,*bs*_ with *y*_R,*bsi*|R,*bs*_, for all *b* ∈ 𝒫, *si* ∈ Ω.

#### A.2.2 Fitness of an average individual

We can obtain the average individual fitness (49)-(50) by taking an average over mutant and resident individual fitness’s (47) and (48). Here, instead, we will provide an alternative derivation that is similar to the the derivation for resident fitness (16): the average individual growth-rate (49)-(50) can be calculated from the mutant-resident group and metacommunity dynamics (42) as

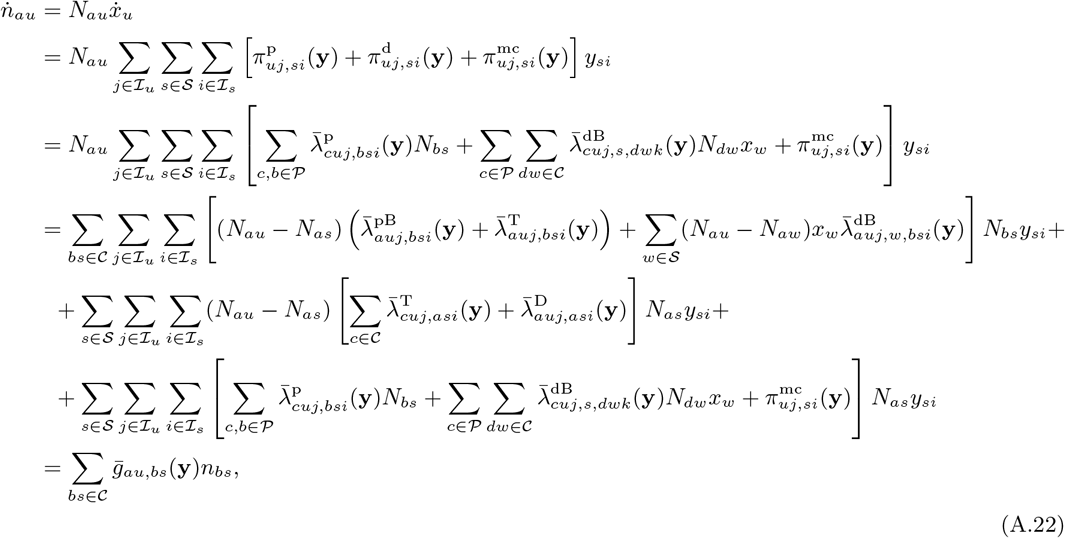

where

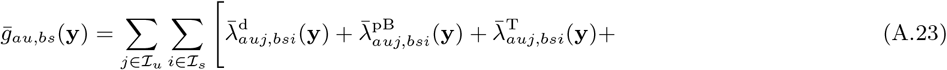

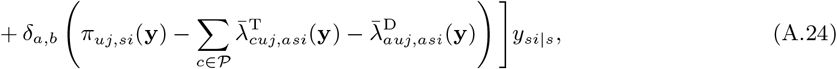

which is the partitioning (50) from the main text.

## B Appendix: Relatedness and genetic correlations

### B.1 Jump-process approach

To facilitate the derivation of the recursion (30) for relatedness (Section 3.4.1), we represent here the metacommunity dynamics as a discret time jump-process. To this end, we focus on a single group and take a probabilistic perspective on the group dynamics by viewing the vector **x**(*t*) := **x** as the probability distribution of the state of some focal group at time *t* (Section 3). The state of the focal group is considered as a random variable which we denote with *S*(*t*) for all *t*. In the continuous time formulation discussed in the main text, the focal group (as well as all other groups in the metacommunity) can thus be seen as undergoing a continuous time (non-homogeneous) Markov chain {*S*(*t*) | *t* ∈ [*t*_0_, *t*_*f*_]} on the state space 𝒮 for some initial time *t*_0_ and final time *t*_*f*_ (and recall that the non-homogeneity is implicit and comes via the group transition matrix (10) being a function of **x**(*t*)). Now, because we want to consider a jump chain (or embedded Markov chain, e.g., Iosifescu, 2007) associated to the above continuous time Markov chain, we are interested in the probabilities at which the i-processes happen. Let Π_*k*_ = *S*(*T*_*k*_) denote the random variable of this jump process giving the state of the group at the *k*th jump starting at time *t*_0_, where each jump time *T*_*k*_ is (also) a random variable. The resident group dynamics of this discrete time Markov chain can then be written as

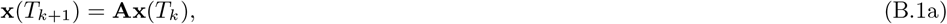

where each element

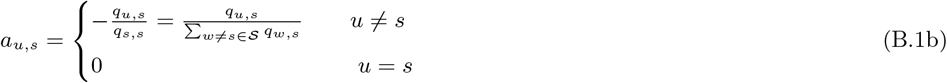

gives the probability at which a group in state *s* ∈ 𝒮 transitions to state *u* ∈ 𝒮 (because no state is absorbing). Using the discrete time group dynamics (B.1) we obtain a jump-process for relatedness (30)-(32) as given in the main text.

### B.2 Mutant-pair dynamics

Here we derive the mutant-pair dynamics (79)-(80) when *δ* = 0 (Section 5.3.3). The ODE for the total (or average) density of local pairs *n*_*abu*_ for all *abu* ∈ ℬ is given, with a slight abuse of notation, identically to (21). By defining *n*_MM,*abs*_ = ∑_*i*∈ℐ_ *i*_*as*_*i*_*bs*_*y*_*si*_ as the density of mutant-pairs *abs* ∈ ℬ, the ODE for mutant-pair dynamics is given as

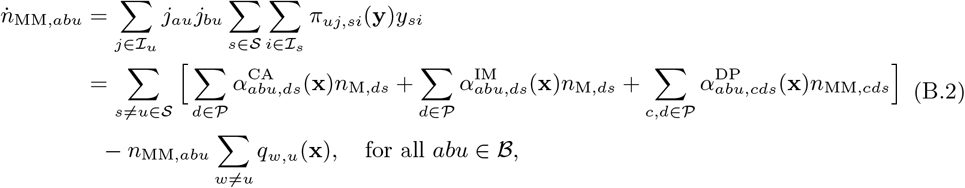

where the above rates are given in terms of i-processes as in (23)-(25) and where we used the consistency relation (58).

We obtain the ODE (79) for the mutant-pair frequencies **p**_MM_ defined in (76) by differentiation and using (21) and (B.2), whereby

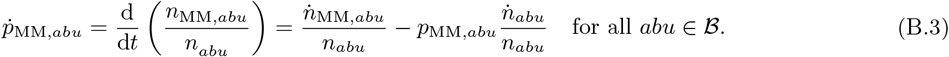

## Notes

### Competing Interest Statement

The authors have declared no competing interest.

